# Seizure-induced Transient Disruptive Changes in Brain Microstructure

**DOI:** 10.1101/2025.05.05.652221

**Authors:** Akihiro Takamiya, Frank Riemer, Vera Jane Erchinger, Max Korbmacher, Hauke Bartsch, Olga Therese Ousdal, Ivan Maximov, Anders Dale, Louise Emsell, Ketil Joachim Oedegaard, Ute Kessler, Leif Oltedal

## Abstract

A leading hypothesis for the mechanisms of electroconvulsive therapy (ECT) posits an initial disruptive effect followed by enhanced neuroplasticity. However, direct evidence of the presumed early disruptive changes, potentially driven by seizures, remains limited. This study examined longitudinal changes in multishell diffusion MRI-derived metrics in 25 individuals with depression, with scans acquired two hours before and after their first ECT session. Follow-up scans were collected within 14 days and 6 months after the last ECT session. To control for potential confounding effects of anesthesia and repeated measurements, two additional groups were included: 16 individuals undergoing short-acting anesthesia and 16 healthy controls without interventions. A multicompartment model was applied to explore extracellular free water and microstructures in the intracellular/extracellular compartments. Whole-brain voxel-wise analyses identified a pattern consistent with vasogenic edema (e.g., increased extracellular free water) in widespread bilateral brain regions following a single ECT session. Increases in brain tissue free water were significantly correlated with electrical stimulus charge (r=0.51, p=0.01) but not with post-ictal reorientation time (r=0.11, p=0.92). These changes were not observed in either control group. Follow-up assessments confirmed that the alterations in tissue free water resolved within 14 days. A single ECT-induced seizure induces transient, reversible changes consistent with vasogenic edema, rather than irreversible cellular injury typically associated with cytotoxic edema. These reversible changes may represent an initial disruptive phase that facilitates subsequent adaptive brain responses, including neuroplasticity and network reorganization underlying the therapeutic effects of ECT.

## Introduction

A seizure is a sudden burst of synchronized electrical activity that disrupts neuronal information flow. While the potential for permanent brain damage remains debated, magnetic resonance imaging (MRI)-detectable brain alterations following seizures are often attributed to disruptive processes such as brain edema. Brain edema, defined as increased water content, is categorized into vasogenic, cytotoxic, and interstitial types (1). Cytotoxic or cellular edema arises from cellular injury (e.g., ischemia or hypoxia) and involves water shifting from extracellular to intracellular spaces. Vasogenic edema results from blood-brain barrier (BBB) disruption, increasing extracellular free water. Interstitial edema occurs when cerebrospinal fluid (CSF) leaks from the ventricles into brain tissue due to elevated intraventricular pressure. Research on epilepsy has reported both cytotoxic and vasogenic edema, depending on seizure type and assessment timing (2, 3). A major challenge in MRI research on spontaneous seizures is distinguishing seizure-induced changes from underlying epileptogenic pathologies (4). Longitudinal MRI studies investigating single seizures are further limited by their unpredictability of seizure onset and the necessity of post-seizure treatments, leaving their direct effects on the human brain largely unknown.

Seizures are not only pathological phenomena but also a key component of electroconvulsive therapy (ECT). ECT is a well-established treatment for various psychiatric symptoms, including depression, mania, and psychosis (5–8), and is also used for pharmacotherapy-resistant psychiatric and motor symptoms in Parkinson’s disease (9). A single ECT session involves electrical stimulation via the scalp under general anesthesia to induce a brief generalized seizure, with multiple sessions typically required to achieve therapeutic effects. One proposed mechanism of ECT action suggests that it temporarily disrupts dysfunctional information flow, followed by enhanced neuroplasticity and network reorganization (10). Since psychiatric symptoms are associated with altered brain networks (11–13), this transient disruption may be necessary for therapeutic efficacy. Over the past decade, accumulating evidence has supported neuroplastic changes following multiple ECT sessions, including transient increases in gray matter volume, particularly in the hippocampus and amygdala (14–18), possibly driven by electrical stimulation (19, 20) or repeated seizures (21). Animal studies have shown increased neurogenesis (22, 23) and synaptic density, the latter of which has been identified as one of the contributors to MRI-detectable hippocampal volume increases following repeated electrically induced seizures (24). While the effects of multiple ECT sessions are well-documented, the impact of a single ECT session on the human brain remains unclear. Investigating this effect not only provides a unique opportunity to study the neurobiological impact of a single seizure but also advances our understanding of ECT mechanisms.

Preclinical animal studies have shown BBB disruption and mild vasogenic edema following electrically induced seizures, partly which may be induced by acute hypertensive effects (25). Human MRI studies have reported mixed findings: one study involving six subjects found increased T2 relaxation time—a marker of edema—two hours after the second ECT session (26), while another study with 15 subjects failed to replicate this result (27). These discrepancies may be due to methodological limitations, including the insensitivity of T2 relaxation time. Recent advances in diffusion MRI enable detection of subtle changes in water diffusivity. Diffusion tensor imaging (DTI), a conventional diffusion MRI modeling approach, provides insight into seizure-related brain edema, with mean diffusivity decreasing in cytotoxic edema and increasing in vasogenic edema (2). Restriction spectrum imaging (RSI)—an advanced diffusion MRI technique—offers deeper insight into intra-voxel microstructural features undetectable by DTI. RSI decomposes diffusion-weighted signals into three compartments: free water (modeled by isotropic diffusion, reflecting water content such as CSF within each voxel), intracellular (modeled by restricted diffusion, reflecting neurite density, axonal myelination, or thin glial processes), and extracellular (modeled by hindered diffusion, representing cellular components in the extracellular space) (28–31). Although a previous cross-sectional RSI study examined temporal lobe epilepsy (29), the findings reflect chronic seizure effects and underlying pathology, rather than the acute impact of a single seizure. Therefore, a well-designed longitudinal RSI study is required to isolate and rigorously evaluate the effects of a single seizure on human brain microstructure.

The present study employs RSI to analyze longitudinal diffusion MRI data acquired two hours before and after a single ECT session to examine the acute effects of a single seizure. Two control groups were included: healthy controls (HC) with no interventions and individuals undergoing short-acting general anesthesia for electrical cardioversion (ECV). These groups were included to control for potential effects of repeated MRI measurements and anesthesia on diffusion MRI-metrics. In addition, we investigated associations between seizure-induced microstructural changes and ECT parameters (e.g., stimulus charge, seizure duration), as well as clinical measures (e.g., post-ictal reorientation time). Finally, we examined the acute effects of a single seizure on ventricular volume as a proxy for intracranial pressure.

## Subjects and Methods

This longitudinal study was approved by the Regional Committee for Medical and Health Research Ethics, REC South―East, Norway (2013/1032) and conducted in accordance with the Declaration of Helsinki. Written informed consent was obtained from all participants. The overall study design was published previously (30), with detailed methods in the Supplementary Material.

### Participants

Participants were recruited from patients referred for ECT at Haukeland University Hospital, Bergen, Norway (September 2013–September 2018). Inclusion criteria were ICD-10-defined unipolar or bipolar depressive disorder, a moderate-to-severe (Montgomery and Asberg Depression Scale >25) depressive episode, and age >18 years. Exclusion criteria included ECT within 12 months, pregnancy, inability to provide consent, and contraindications to MRI. Two control groups were recruited: HC and individuals receiving general anesthesia for ECV for atrial fibrillation to control for repeated MRI measurements and anesthesia effects, respectively.

### ECT

ECT was administered three times a week using a Thymatron System IV, delivering brief pulse (0.5 ms) right unilateral (RUL) stimulation. The initial stimulus charge determined by age (30). Thiopental or propofol was used for anesthesia, and succinylcholine for muscle relaxation. Seizure duration was monitored via two-channel EEG (Fp1-M1, Fp2-M2, international 10-20 system).

### Image acquisition and processing

Image acquisition is presented in Supplementary Material. MRI scans were conducted two hours before (timepoint 1: TP1) and after (TP2) the first ECT session. Participants in the HC and ECV groups underwent MRI at identical timepoints (8 AM and 12 PM). T1-weighted images were processed using the longitudinal pipeline of the Computational Anatomy Toolbox (CAT12, version 12.9) for ventricular/CSF volume analysis, brain mask creation, and registration parameter refinement for diffusion-weighted imaging. Bilateral ventricular volumes were calculated using the Hammers Atlas and normalized by total intracranial volume (TIV). Tissue-specific masks [i.e., gray matter (GM), white matter (WM), or CSF] were created in MNI152NLin2009cAsym space. The cerebellum and lower brain stem, which were not fully covered in all scans, were excluded.

Diffusion-weighted images were visually inspected for artifacts, leading to an exclusion of one participant due to excessive motion artifact. Images were pre-processed using an in-house pipeline, including correction for B0 distortion, motion and eddy current distortions, gradient nonlinearity distortions, and head motion. To enhance registration accuracy, B0 images were registered to skull-stripped T1-weighted images using Advanced Normalization Tools (ANTs: https://github.com/ANTsX/ANTs), and transformation matrices were applied to the pre-processed diffusion-weighted images. To optimize intra-subject alignment, study-specific T1 templates were created using subject-specific average images between the time points with <u>antsMultivariateTemplateConstruction2.sh</u>, and all images were subsequently warped to the study-specific template space (31–34). All images were then warped to the standard MNI152NLin2009cAsym space and smoothed with a 2-mm full-width at half-maximum (FWHM) Gaussian kernel.

The RSI model was fitted using an in-house Matlab script (29, 35). Theoretical background and technical details are presented elsewhere (28). RSI-derived metrics (i.e., extracellular free water, intracellular restricted diffusion, and extracellular hindered diffusion) were normalized by taking the norm (square root of the sum of squares) of each fractional map and dividing by the norm of all fractional maps, making them unitless with range between 0 to 1. In an additional analysis, mean diffusivity maps were generated using diffusion tensor imaging with b=1000 in FSL (Supplementary Figure 1).

To investigate short- and long-term RSI changes, follow-up MRI data were collected at 7–14 days post-ECT (TP 3, n=23) and 6 months post-ECT (TP 4, n=17). Because the primary aim of this study was to investigate the effects of a single seizure on the human brain, this follow-up analysis was limited to the RSI changes identified in the whole-brain analysis between TP1 and TP2 to determine whether these changes were transient.

### Statistical analysis

To investigate the acute effects of a single ECT session on brain microstructure, voxel-wise paired t-tests were conducted in the ECT cohort using FSL’s “randomize” tool with 5,000 permutations. The statistical significance threshold was set at family-wise error (FWE)-corrected p <0.05, determined by threshold-free cluster enhancement (TFCE) (36). As RSI modeling can be applied to both WM and GM (28, 29, 37, 38), primary analyses included both tissue types.

Based on the results of whole-brain analyses in the ECT cohort, three additional analyses were conducted. We calculated median RSI values in identified brain regions to reduce the effects of outliers. First, to explore the effects of ECT parameters, correlation analyses were conducted between changes in RSI metrics and stimulus charge or seizure duration. Second, post-ictal reorientation time, a predictor of retrograde amnesia (39), was also examined to assess its association with RSI changes. Third, to ensure that the observed regional changes were not driven by time effects or anesthesia, paired t-tests were conducted in the HC and ECV groups using masks derived from the ECT cohort results. For interpretability, boxplots of RSI values were created for the three groups. Fourth, to explore short- and long-term trajectories of the identified RSI changes, linear mixed--effects models were employed, with time included as a fixed effect, subject as a random effect, and age and sex as covariates, to account for missing values at TP3 and TP4. The statistical significance level was set at p <0.05/the number of analyses for correlations and the linear mixed effects model, and FWE-corrected p <0.05 for voxel-wise analysis.

To assess the impact of a single seizure on CSF/ventricular volume changes, deformation-based morphometry (DBM) was conducted. DBM is a voxel-wise method for detecting macroscopic morphological changes by utilizing image registration parameters rather than tissue segmentation (40). Using the CAT12, normalized Jacobian maps from each time point were extracted and smoothed with a 2 mm FWHM Gaussian kernel. Voxel-wise paired t-tests were conducted using FSL’s “randomize” tool with 5000 permutations, applying an FWE-corrected significance threshold of p <0.05. An additional region-of-interest (ROI) analysis was conducted using the Hammers Atlas to specifically investigate changes in temporal horn volume, a region sensitive to intracranial pressure fluctuations. To assess volumetric changes in CSF spaces, linear mixed-effects models were applied to TIV-normalized volumes of the temporal horn, lateral ventricles excluding temporal horn, and third ventricle. Age and sex were included as covariates, and group (ECT, HC, ECV) by time (TP1, TP2) interactions was tested, with statistical significance set at p<0.05.

## Results

We analyzed 154 MRI scans from 57 participants (25 ECT, 16 HC, and 16 ECV). There were significant group differences in age (F_2,31.1_=29.7, p<0.001) and female proportion (Fisher’s exact test: p<0.001). Post-hoc pairwise analyses revealed that the mean age in the ECV group [mean (SD) = 65.5 (6.5)] was higher than the ECT group [44.6 (15.3)] and HC [40.1 (15.9): both p <0.001], while there was no significant difference in the mean age between the ECT and the HC group (p=0.64). The female proportion was significantly lower in the ECV group (1/16) compared to the ECT (12/25: Fisher’s exact test: p=0.006) and HC group (5/16: Fisher’s exact test: p<0.001), while there was no significant difference in female proportion between ECT group and HC (Fisher’s exact test: p=0.22). Other clinical characteristics of ECT cohort are presented in Table 1.

**Table 1.** Clinical characteristics of the participants.

|  | ECT | HC | ECV | Statistical<br>value | p-value |
| --- | --- | --- | --- | --- | --- |
| n | 25 | 16 | 16 |  |  |
| age, years | 44.6 (15.3) | 40.1 (15.9) | 65.5 (6.5) | $F_{2,31.1}=29.7$ | <0.001 |
| female, % | 48.0 (12/25) | 68.8 (11/16) | 6.3 (1/16) |  | <0.001 |
| MDD/Bipolar | 21/4 |  |  |  |  |
| Psychotic, % | 28.0 (7/25) |  |  |  |  |
| Melancholic, % | 72.0 (18/25) |  |  |  |  |
| Family<br>history, % | 64.0% (16/25) |  |  |  |  |

Abbreviation: ECT, electroconvulsive therapy; HC, healthy control; ECV, electrical cardioversion; MDD, major depressive disorder

### Acute effects of a single seizure on brain microstructure

Voxel-wise analyses identified increased free water and decreased restricted diffusion two hours after the first ECT session, with no significant changes in hindered diffusion (Figure 1A, 1B). The identified regions were primarily located in periventricular regions and subcortical areas surrounding the hippocampus, with minimal involvement of cortical regions (Figure 1C, 1D). The result of mean diffusivity is reported in Supplementary Figure 1 and effect size maps of RSI- and DTI-metrics are presented in Supplementary Figure 2. Separate analyses of GM and WM are reported in Supplementary Figure 3, 4.

**Figure 1.**
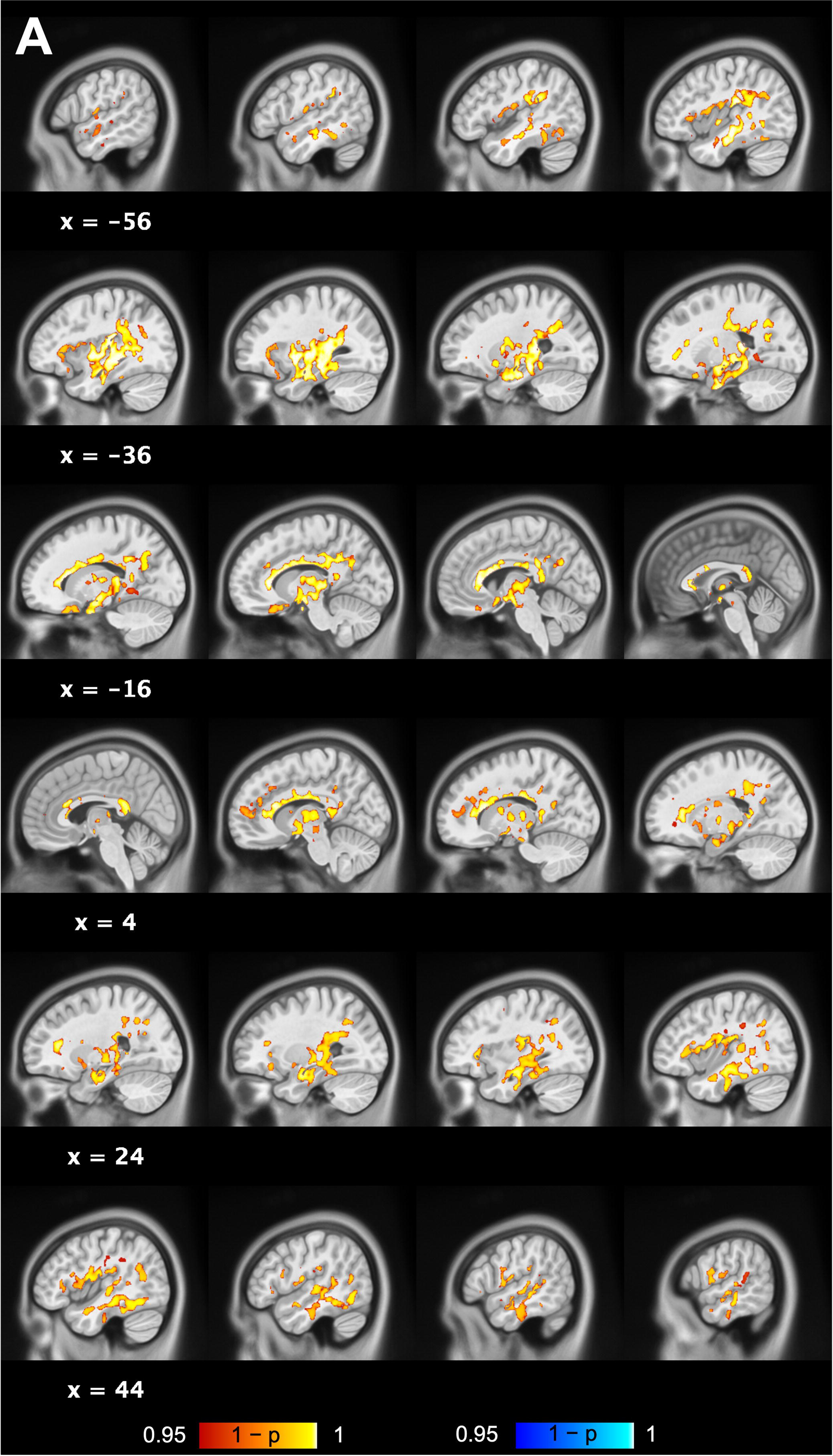
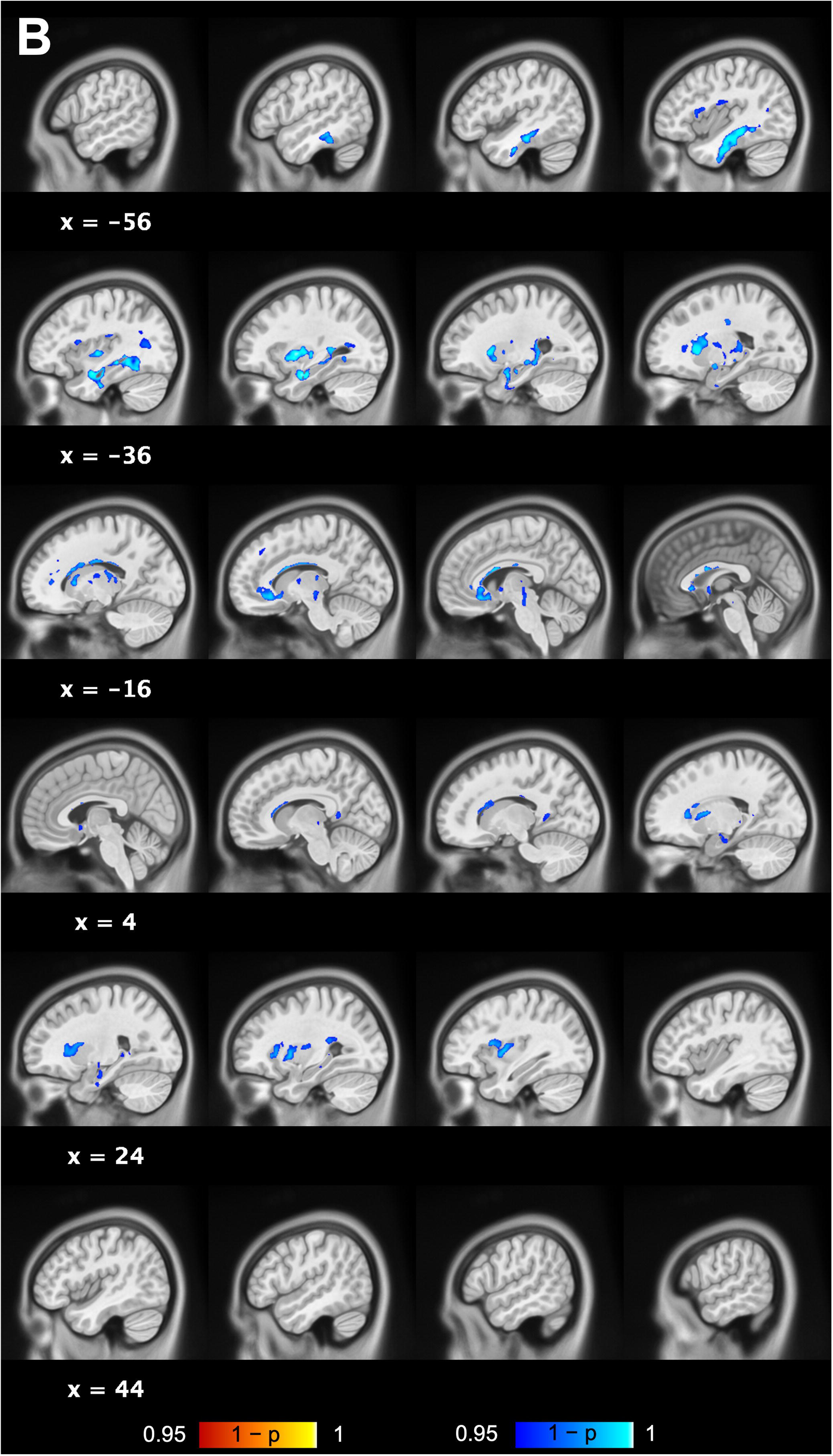
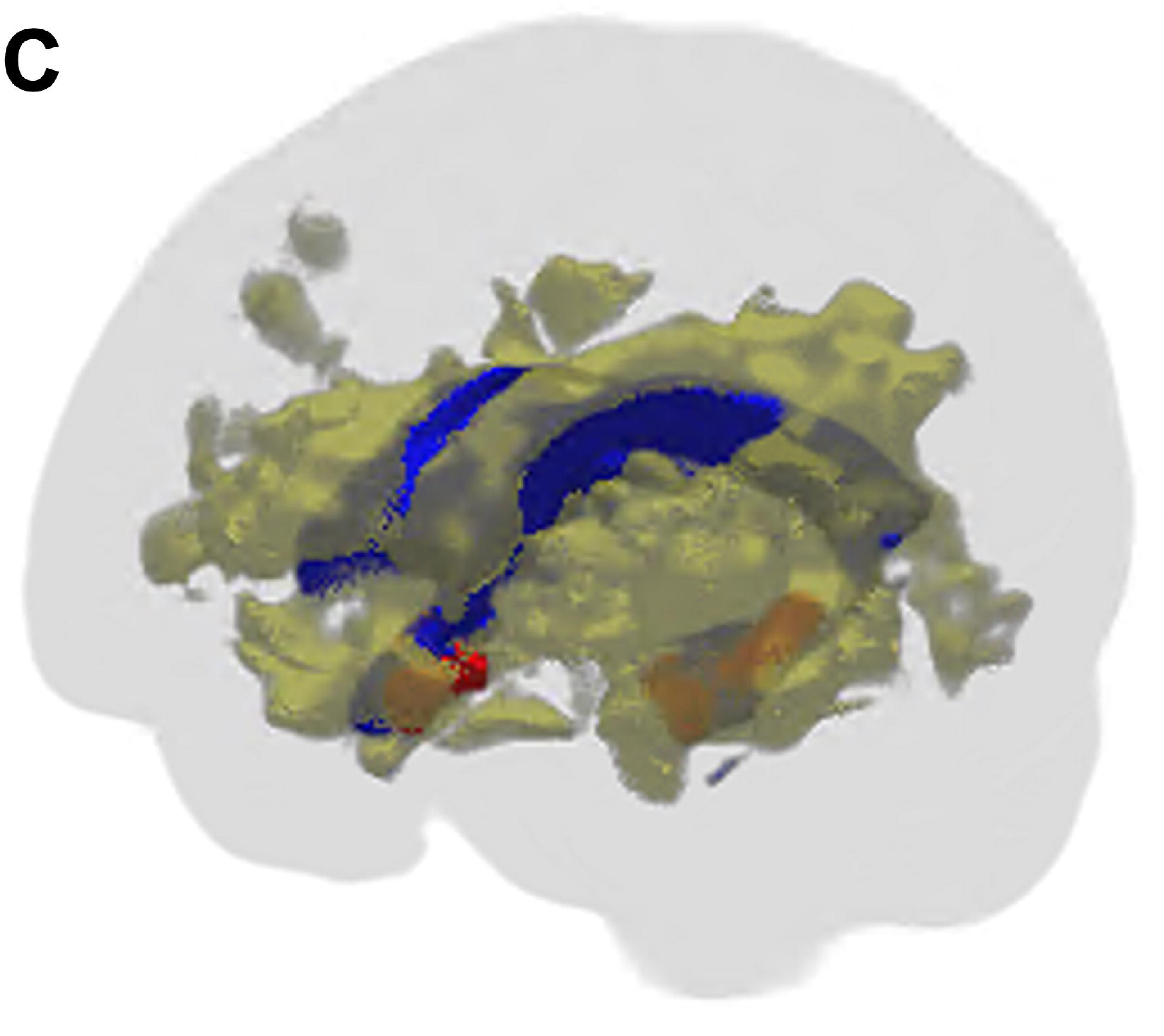
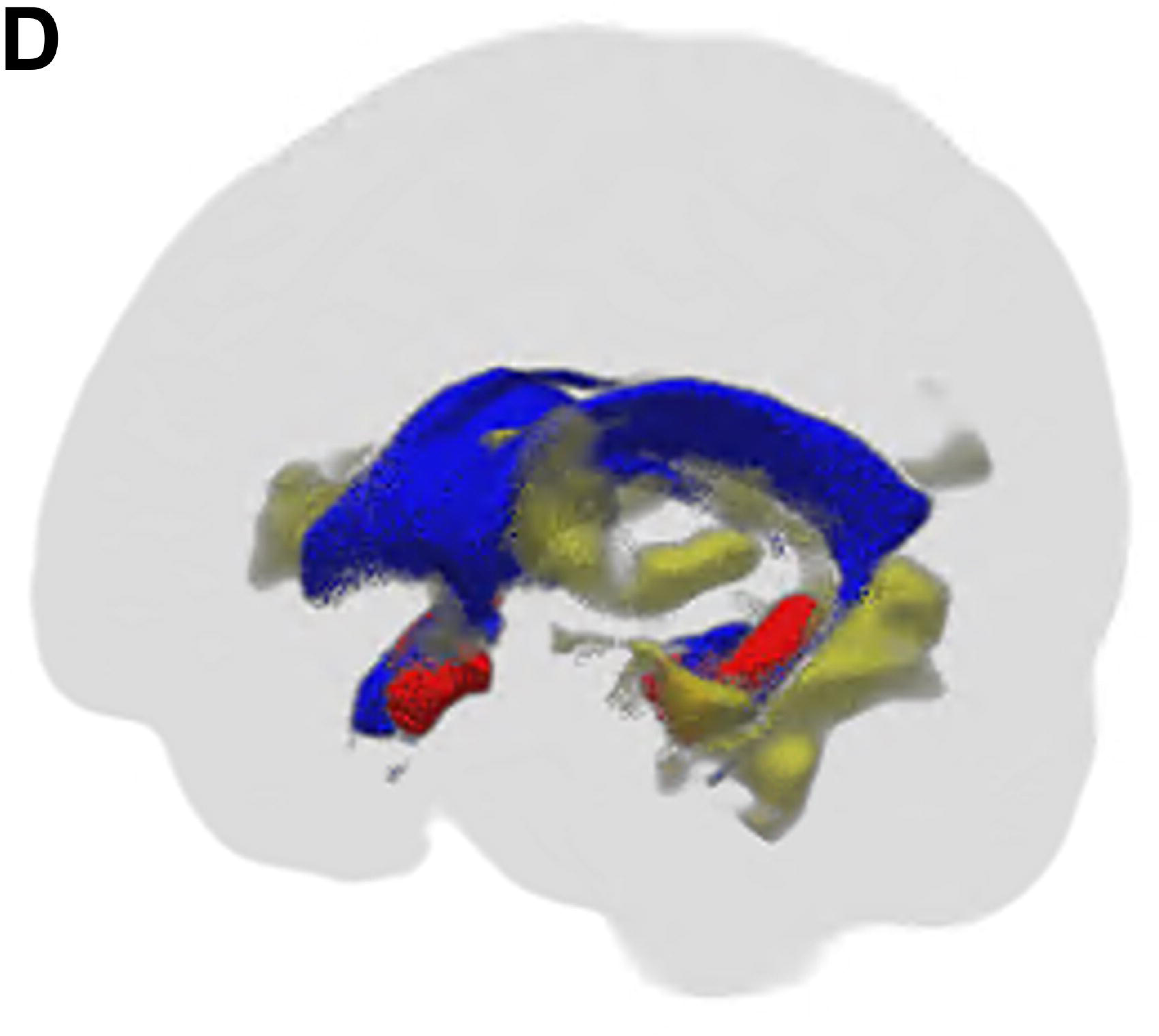
Results of the whole-brain voxel-wise analysis of RSI-metrics. The voxel-wise analyses identified increased free water (**A**) and decreased restricted diffusion (**B**) two hours after a single seizure. Red indicated increases, while blue indicates decreases in each metric following a single seizure. The statistical significance level was set at FWE-corrected p <0.05 determined by TFCE. 3D representation of the results of free water (**C**) and restricted diffusion (**D**). The identified regions (yellow) are also presented on the 3D glass brain with the lateral ventricles (blue) and the hippocampus (red) for interpretation purposes.

### Association of stimulus charge with brain microstructural changes

Stimulus charge correlated with free water changes (r=0.51, df=23, p=0.0098; <0.05/4=0.0125; Figure 2), but not with restricted diffusion (r=0.16, df=23, p=0.44). Seizure duration was not associated with changes in free water (r=–0.30, df=23, p=0.15) or restricted diffusion (r=–0.13, df=23, p=0.22).

**Figure 2.**
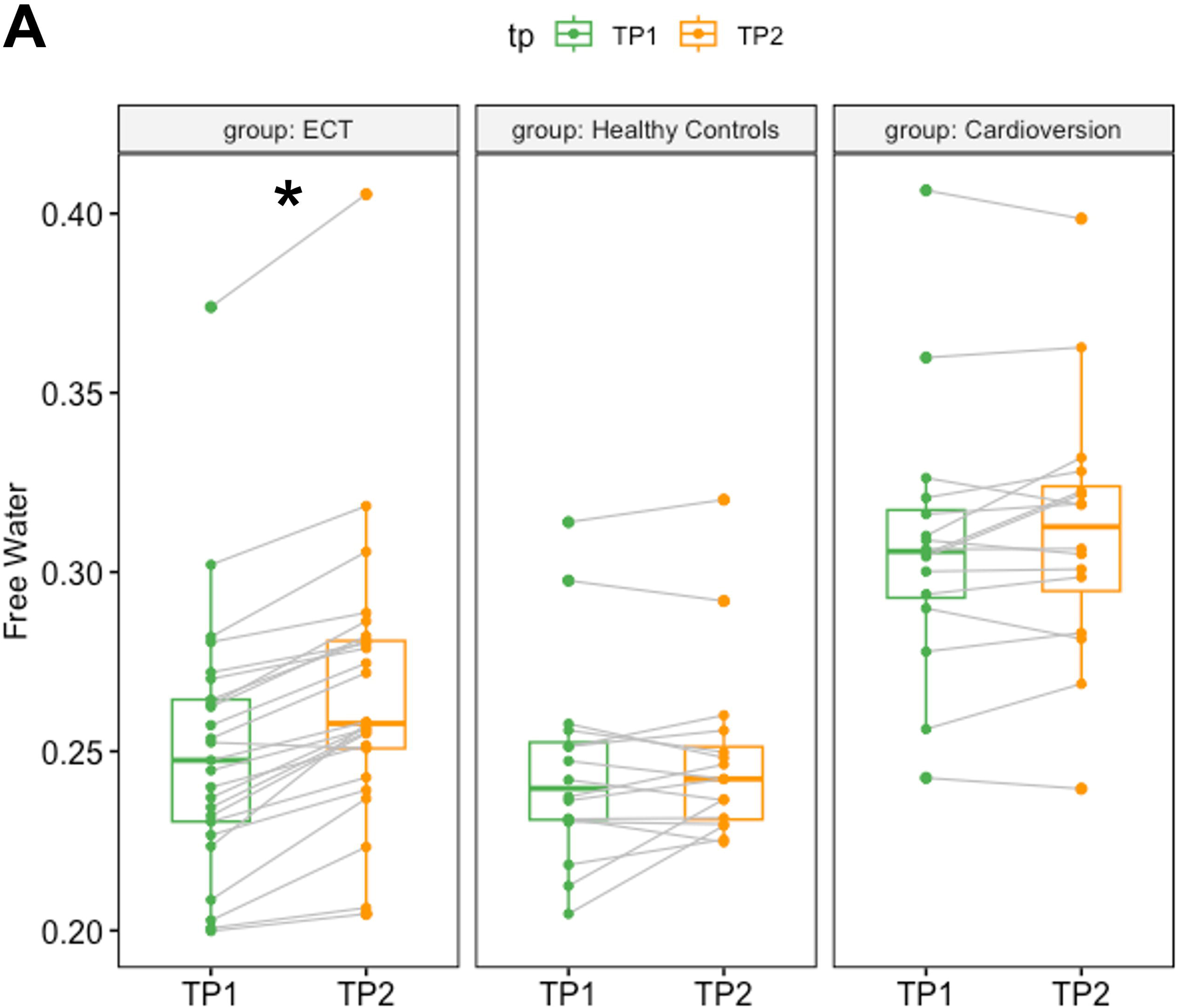
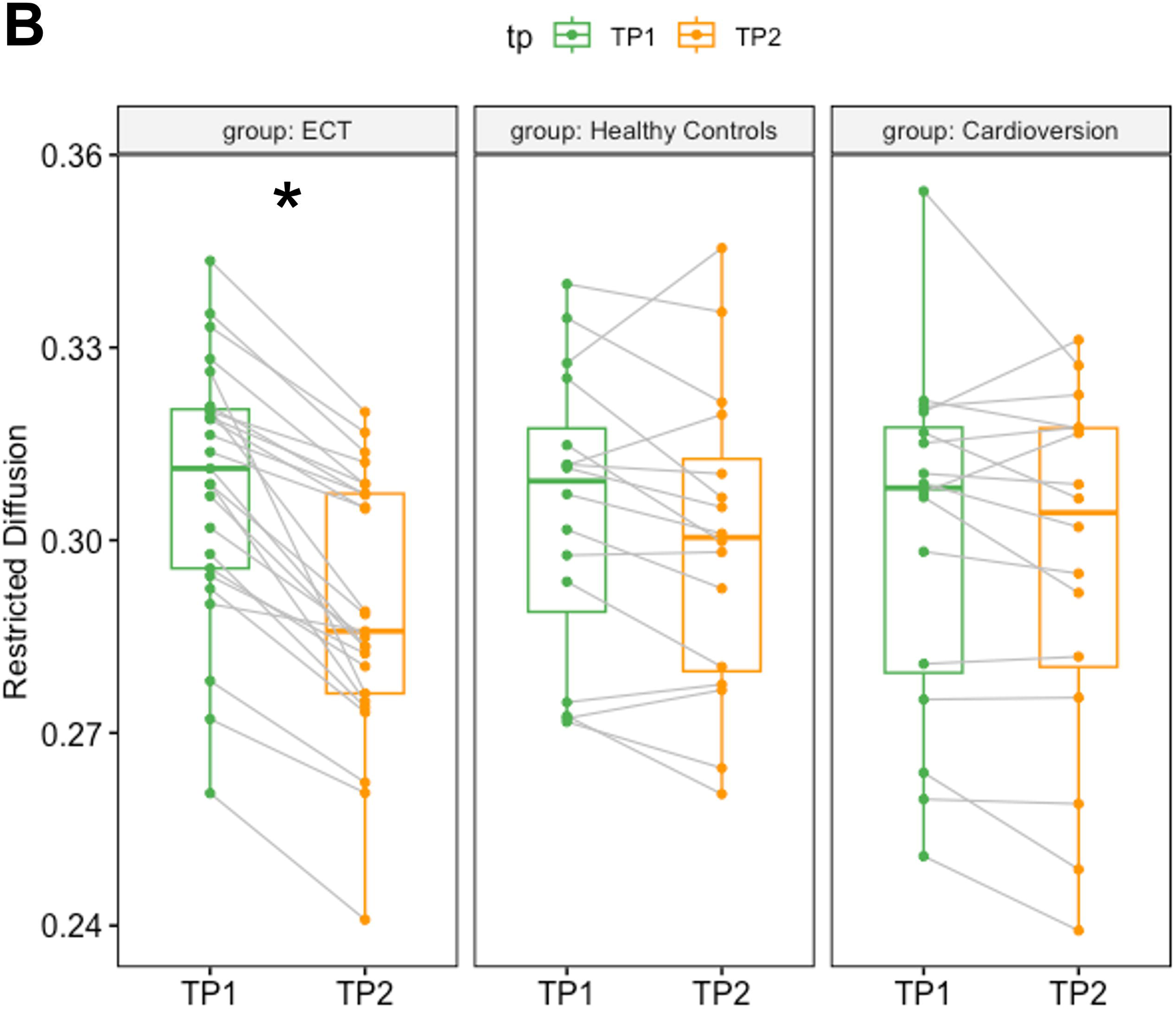
Association between ECT-related parameters and changes in the RSI-metrics. Changes in free water was significantly correlated with stimulus charge (**A**: r=0.51, df=23, p=0.0098), but not with seizure duration (**B**: r=–0.30, df=23, p=0.15). Changes in restricted diffusion was not associated with stimulus charge (**C**) or seizure duration (**D**).

### Association of post-ictal reorientation time with brain microstructural changes

No significant correlations were observed between post-ictal reorientation time and changes in free water (r=0.11, df=23, p=0.92) or restricted diffusion (r=0.10, df=23, p=0.64) (Supplementary Figure 5).

### Effects of time and anesthesia

Voxel-wise paired t-tests between the two timepoints in the HC and ECV groups showed no significant changes in free water or restricted diffusion. Results remained unchanged at a more liberal threshold (FWE-corrected p <0.1) to account for smaller sample sizes in control groups. Boxplots of extracted values for the three groups are presented in Figure 3.

**Figure 3.**
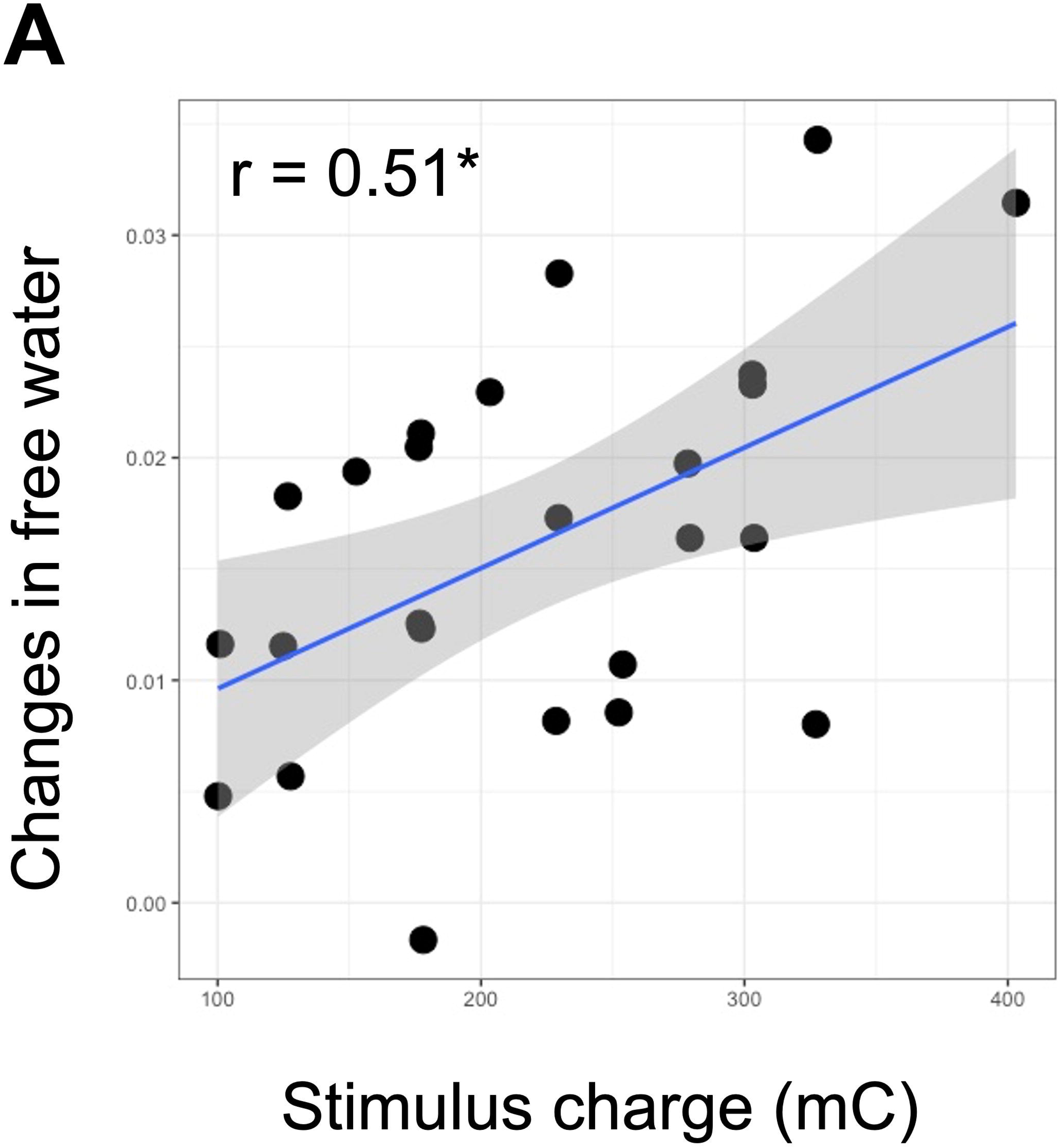
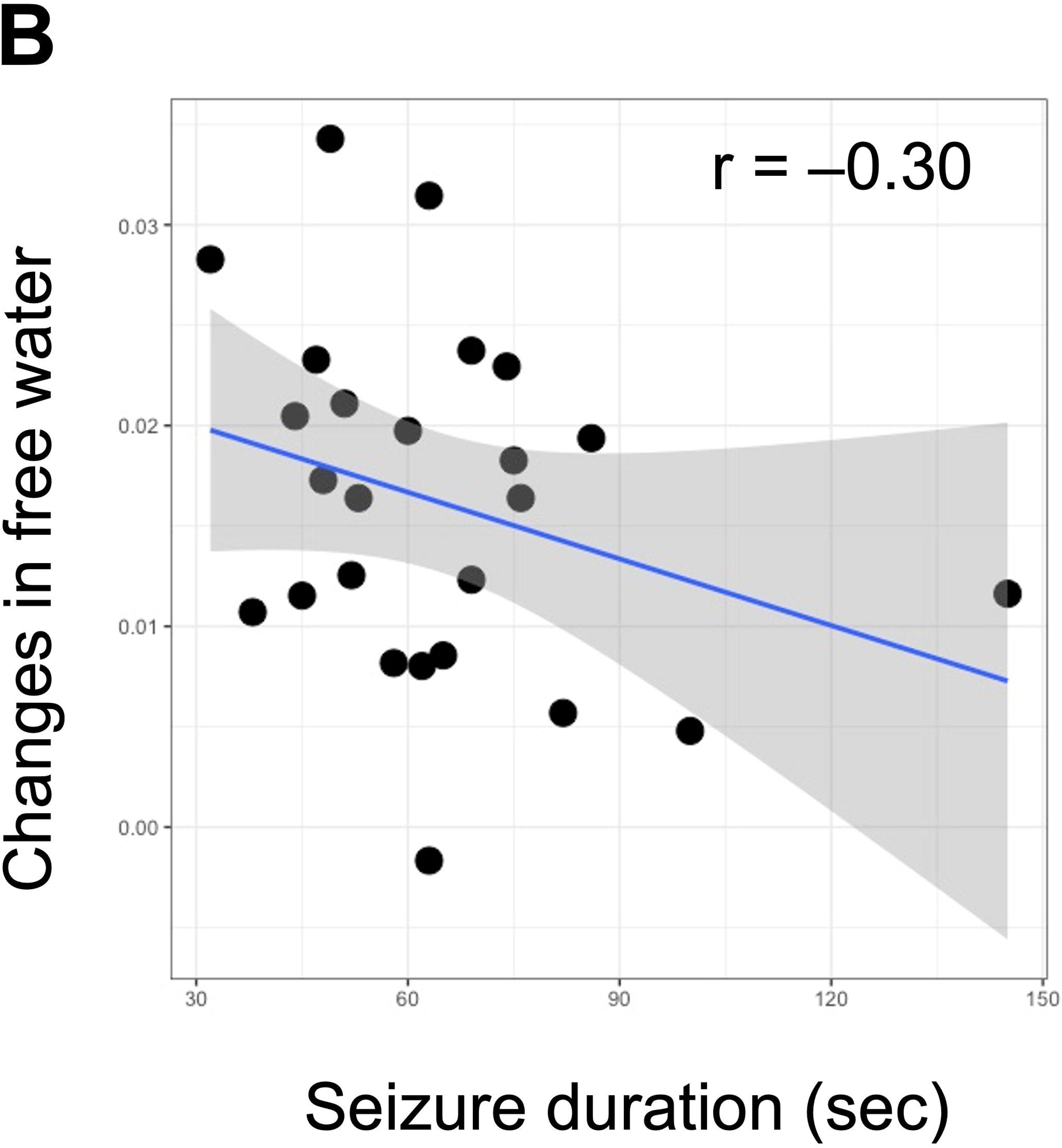
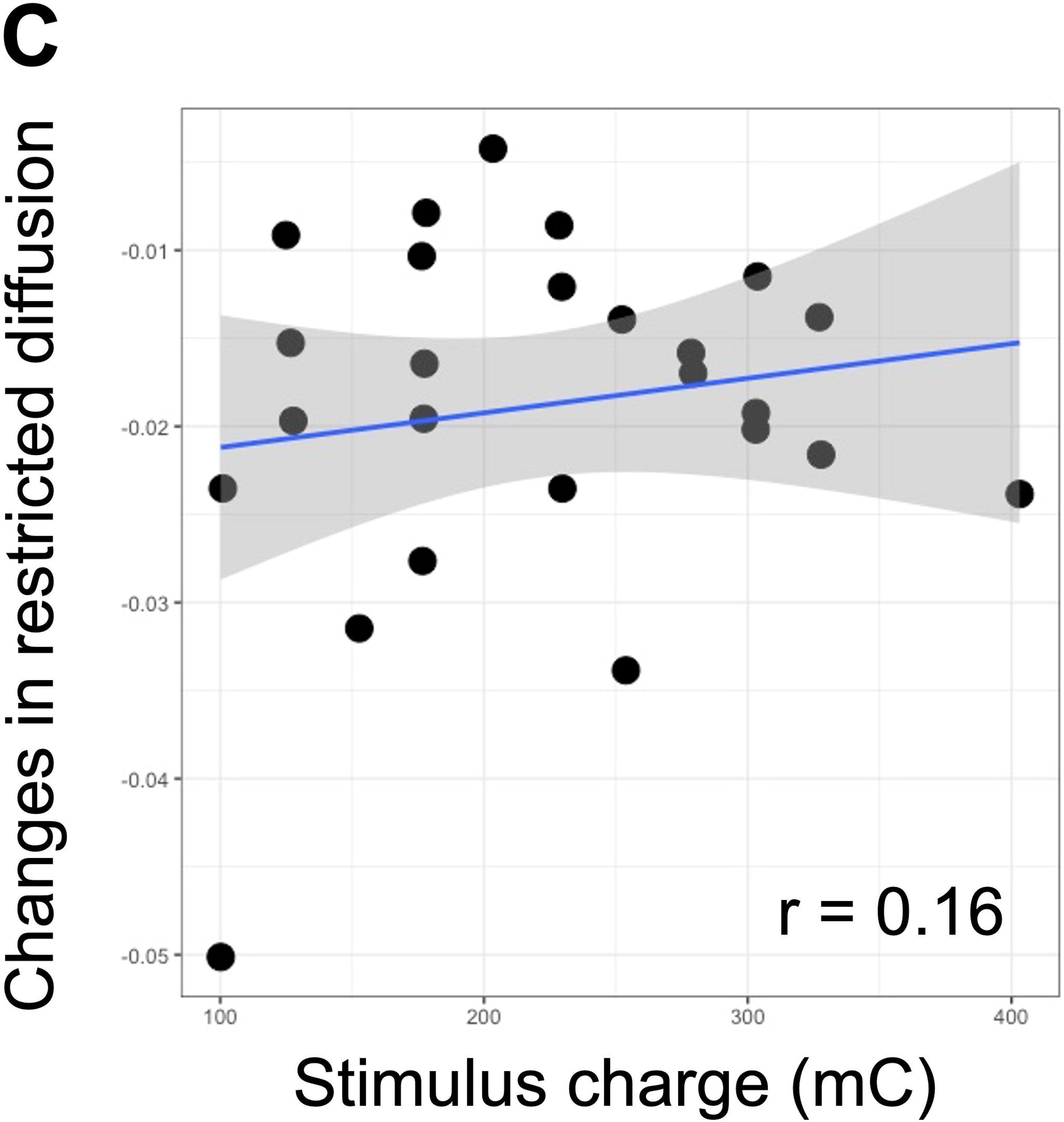
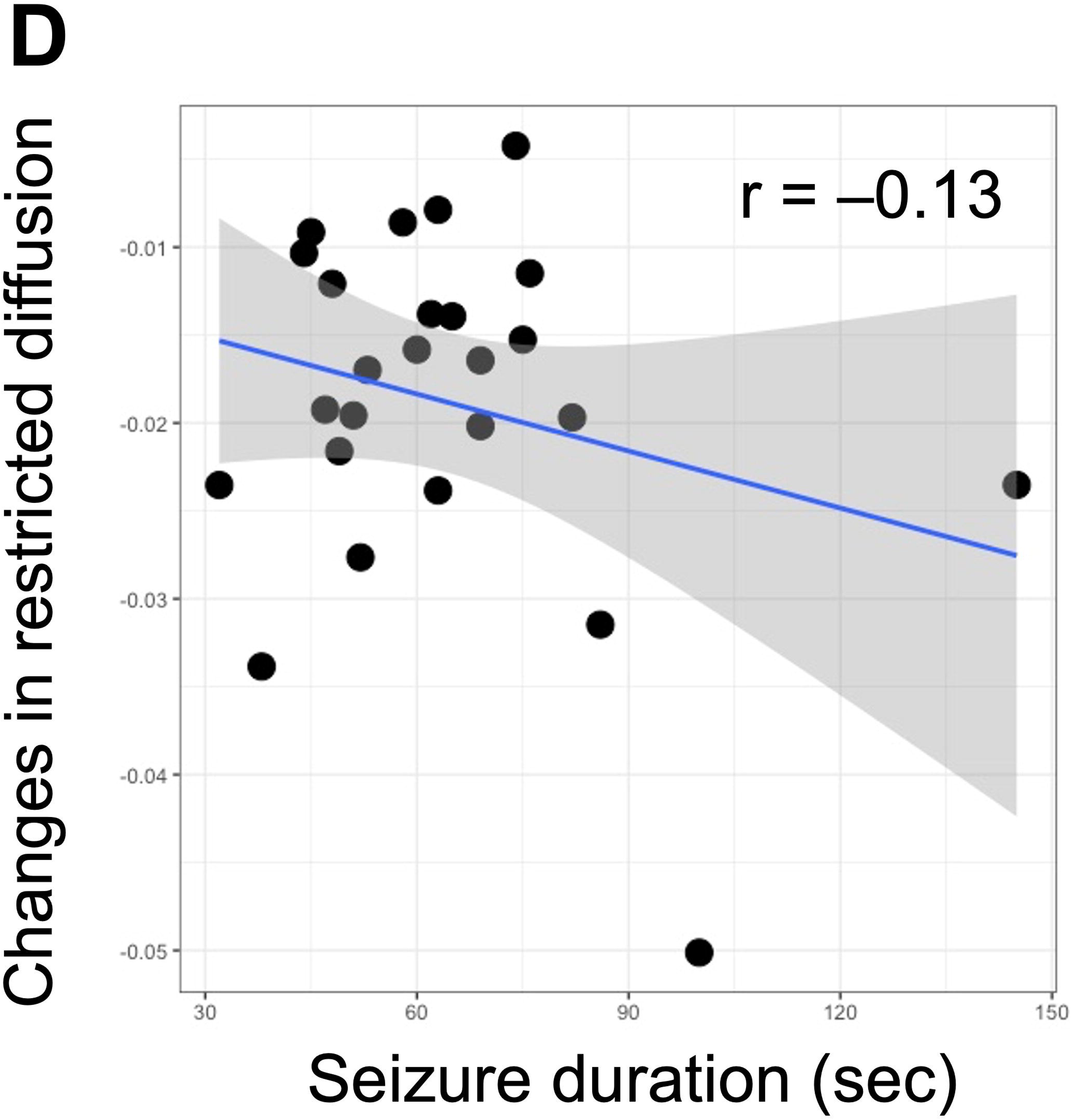
No significant increases in free water or decreases in restricted diffusion in the HC or ECV group. Extracted values of free water (**A**) and restricted diffusion (**B**) in the regions identified in the ECT group showed no significant changes in the HC group (free water: t=1.7, p=0.12; restricted diffusion: t=–1.8, p=0.09) or the ECV group (free water: t=1.3, p=0.23; restricted diffusion: t=–1.9, p=0.08). These boxplots are presented for illustrative purposes. The voxel-wise paired t-tests in the HC or ECV groups did not show significant results. Statistics on pre-post changes in the ECT cohort are not presented to avoid analytical circularity, as the values were extracted from the regions identified as statistically significant in a voxel-level paired t-test within the same cohort (Figure 1). TP, timepoint: TP1 is around 8:00 am and TP2 is around 12:00 pm in the same day.

### Long-term changes in RSI-metrics

Linear mixed effects models with post-hoc pairwise comparisons revealed that increased free water and decreased restricted diffusion observed at TP2 returned to baseline levels by TP3 and TP4, respectively (Supplementary Figure 6), suggesting that the observed changes following a single seizure was transient and reversible.

### Acute effects of a single seizure on ventricular volume

DBM revealed significant CSF space expansion in the lateral ventricles, sylvian fissure, and choroidal fissure bilaterally. No significant changes were observed in the HC or ECV groups. ROI analysis confirmed increased volumes in the temporal horn of the lateral ventricles. All results of the DBM and ROI analyses are presented in Supplementary Figure 7.

## Discussion

This study provides several key findings: 1) a single seizure induced by electrical stimulation increased free water in brain tissue; 2) the degree of free water increase correlated with stimulus charge; 3) the observed changes were not attributable to repeated measurements or anesthesia; 4) free water returned to baseline levels by 7–14 days post-ECT, and 5) ventricular enlargement was detected following a single seizure, indicative of increased intracranial pressure. These findings contribute to understanding seizure-induced neurobiological changes and indicate that the effects are consistent with vasogenic and/or interstitial edema, rather than cytotoxic edema. The transient and reversible nature of these changes suggests a low likelihood of irreversible cellular injury. Furthermore, the observed transient edematous changes are consistent with the initial disruptive phase proposed in the “disruption, potentiation, rewire” hypothesis for the mechanism of ECT action (10).

Previous research on epilepsy has reported both cytotoxic and vasogenic edema following seizures, depending on the timing of MRI assessments, seizure type, and severity (2). Cytotoxic edema is characterized by a shift of water into the intracellular space, typically manifesting as reduced mean diffusivity, whereas vasogenic edema is characterized by extracellular water accumulation, presenting as increased free water and mean diffusivity (1, 41). Our findings of increased free water and mean diffusivity suggest that vasogenic edema predominates two hours after a single seizure. The observed reduction in intracellular restricted diffusion further supports this interpretation. Unlike pathological epileptic seizures, which are often associated with cytotoxic edema, the absence of cytotoxic edema in our study may be due to the controlled nature of ECT-induced brief seizures (e.g., mean duration = 64.2 seconds) performed under general anesthesia with adequate oxygenation. This interpretation is supported by animal studies reporting no evidence of cellular death following electrically induced seizures (22, 42), as well as human research showing transient astrocytic reactivity without changes in brain injury markers (e.g., neurofilament light chain) following a single seizure (43). Together, these findings support the notion that seizures do not necessarily cause brain damage and suggest that factors such as seizure severity (e.g., status epilepticus), hypoxia, or underlying epileptogenic pathologies may play a critical role in determining the risk of brain injury.

Vasogenic edema results from increased BBB permeability, as observed in both human and animal studies following epileptic (44) and electrically induced seizures (25, 45). Although the mechanisms underlying seizure-related BBB changes remain unclear, neuroinflammation may contribute. Previous studies have consistently demonstrated elevated levels of proinflammatory cytokines (e.g., IL-6) within one (46) to six hours (47) after a single ECT session, correlating with stimulus charge (47). In our study, stimulus charge was positively associated with increased free water (Figure 2A), potentially reflecting elevated BBB permeability and inflammatory responses. However, since stimulus charge was determined by age—with older participants receiving higher electrical doses—this study cannot fully disentangle the effects of age from those of electrical stimulation due to their strong correlation. Notably, older age has been linked to increased BBB permeability in humans, particularly in the hippocampus (48), suggesting that both factors may lead to similar changes (i.e., increased BBB permeability).

Another possible contributor to increased free water is elevated intracranial pressure (25). An animal study demonstrated that seizure-induced increases in BBB permeability were abolished when hypertensive surges were prevented, suggesting that acute hypertensive responses may directly affect BBB integrity (25). Our findings of ventricular enlargement, particularly in the temporal horn (Supplementary Figure 7), support this interpretation. Therefore, interstitial edema due to increased intracranial pressure, in addition to vasogenic edema, may contribute to the observed free water increases, particularly in periventricular regions (Figure 1C).

Although this study aimed to investigate the acute disruptive effects of a single seizure, it does not imply that ECT is harmful. ECT is a highly effective treatment for severe psychiatric disorders, with a well-established safety profile (49, 50), a lower mortality rate than general anesthesia (51), and no increased risk of developing dementia (52, 53). Importantly, ECT has demonstrated antiepileptic properties (54), in contrast to pathological epileptic seizures. Several unique features differentiate ECT from epileptic seizures: controlled electrical stimulation, brief seizure duration, administration under general anesthesia with adequate oxygenation, and repeated induction of seizures over a treatment course. While our findings offer novel insights into seizure-induced MRI changes, the absence of cytotoxic edema highlights a key distinction between ECT and pathological epilepsy, particularly status epilepticus. Moreover, the observed increase in free water was transient and reversible (Supplementary Figure 6), suggesting that a single ECT session induces only temporary disruption, rather than permanent brain damage. This interpretation is supported by the absence of macroscopic radiological evidence of brain injury, including microbleeds (55). Rather than being harmful, such transient and reversible disruptive effects may serve as necessary precursors—temporarily disrupting pathological information flow or abnormal brain states associated with depression—to facilitate subsequent neuroplasticity (14–18) and reorganization of neuronal networks (56–58), which are thought to underlie the therapeutic effects of ECT. This perspective may be in line with the concept of hormesis (59), a biphasic dose-response phenomenon in which low-dose exposures (e.g., well-controlled brief seizures) trigger beneficial adaptations or reparative overcompensatory processes, while high-dose exposures (e.g., pathological status epilepticus) result in damaging effects such as irreversible cellular injury.

A key strength of this study lies in its unique design, which allowed us to examine the acute effects of a single seizure rather than repeated seizures. MRI data acquired two hours before and after the first ECT session enabled precise characterization of seizure-induced changes. The inclusion of two control groups enhanced the validity of our findings, confirming that observed increases in free water and decreases in restricted diffusion were not attributable to repeated measurements or anesthesia. Nevertheless, some limitations should be acknowledged, as they are inherent to the clinical nature of this study. First, no significant association was found between increased free water and post-ictal reorientation time, although the latter is considered a proxy for long-term memory dysfunction (39) rather than a direct indicator of acute cognitive effects. Second, the ECV cohort was older than the other groups, reflecting the clinical demographics of individuals undergoing cardioversion. While unavoidable, we focused on within-subject longitudinal changes to minimize age-related confounds.

### Conclusion

A single seizure induced by electrical stimulation induced acute vasogenic and/or interstitial edema, rather than cytotoxic edema. These results suggest that a single seizure induces reversible disruptive changes and is unlikely to cause irreversible cellular injury.

## Acknowledgements

The study was funded by Western Norway Regional Health Authority grant no 911986 (to KJO) and 912238 (to LO). AT was supported by a postdoctoral fellowship (FWO/1283524N).

The authors would like to thank Leila Marie Frid for data organization.

## Author contributions

Dr Takamiya had full access to all of the data in the study and take responsibility for the integrity of the data and the accuracy of the data analysis.

*Concept and design:* Leif Oltedal, Ketil J Oedegaard, Anders Dale, Renate Grüner, Ute Kessler,

*Acquisition, analysis, or interpretation of data:* Vera Jane Erchinger, Leif Oltedal, Olga Therese Ousdal, Max Korbmacher, Ivan Maximov, Ute Kessler, Frank Riemer

*Drafting of the manuscript:* Akihiro Takamiya

*Critical review of the manuscript for important intellectual content:* All authors

*Statistical analysis:* Akihiro Takamiya, Max Korbmacher

*Obtained funding:* Leif Oltedal and Ketil J Oedegaard

*Supervision:* Leif Oltedal

## Data availability

All data that support the findings of this study are stored in a secured online research platform. Anonymized data are available upon reasonable request and approval by the local Ethics Committee.

## References

1. Hu S, Exner C, Sienel RI, et al. Characterization of Vasogenic and Cytotoxic Brain Edema Formation After Experimental Traumatic Brain Injury by Free Water Diffusion Magnetic Resonance Imaging. J Neurotrauma. 2024;41(3-4):393–406. doi:10.1089/neu.2023.0222

2. Briellmann RS, Wellard RM, Jackson GD. Seizure-associated abnormalities in epilepsy: Evidence from MR imaging. Epilepsia. 2005;46(5):760–766. doi:10.1111/j.1528-1167.2005.47604.x

3. Concha L, Kim H, Bernasconi A, Bernhardt BC, Bernasconi N. Spatial patterns of water diffusion along white matter tracts in temporal lobe epilepsy. Neurology. 2012;79(5):455–462. doi:10.1212/WNL.0b013e31826170b6

4. Mariajoseph FP, Muthusamy S, Amukotuwa S, Seneviratne U. Seizure-induced reversible MRI abnormalities in patients with single seizures: a systematic review. Epileptic Disord. 2021;23(4):552–562. doi:10.1684/epd.2021.1300

5. Espinoza RT, Kellner CH. Electroconvulsive therapy. N Engl J Med. 2022;386:667–672. doi:10.1056/NEJMra2034954

6. Schoeyen HK, Kessler U, Andreassen OA, et al. Treatment-resistant bipolar depression: A randomized controlled trial of electroconvulsive therapy versus algorithm-based pharmacological treatment. Am J Psychiatry. 2015;172(1):41–51. doi:10.1176/appi.ajp.2014.13111517

7. Popiolek K, Bejerot S, Landén M, Nordenskjöld A. Association of Clinical and Demographic Characteristics with Response to Electroconvulsive Therapy in Mania. JAMA Netw Open. 2022;5(6):E2218330. doi:10.1001/jamanetworkopen.2022.18330

8. Sinclair DJM, Zhao S, Qi F, Nyakyoma K, Kwong JSW, Adams CE. Electroconvulsive therapy for treatment-resistant schizophrenia. Cochrane Database Syst Rev. 2019;2019(3). doi:10.1002/14651858.CD011847.pub2

9. Takamiya A, Seki M, Kudo S, et al. Electroconvulsive Therapy for Parkinson’s Disease: A Systematic Review and Meta-Analysis. Mov Disord. 2021;36(1):50–58. doi:10.1002/mds.28335

10. Ousdal OT, Brancati GE, Kessler U, et al. The Neurobiological Effects of Electroconvulsive Therapy Studied Through Magnetic Resonance: What Have We Learned, and Where Do We Go? Biol Psychiatry. 2022;91(6):540–549. doi:10.1016/j.biopsych.2021.05.023

11. Mayberg HS, Liotti M, Brannan SK, et al. Reciprocal limbic-cortical function and negative mood: Converging PET findings in depression and normal sadness. Am J Psychiatry. 1999;156:675–682.

12. Kaiser RH, Andrews-Hanna JR, Wager TD, Pizzagalli DA. Large-scale network dysfunction in major depressive disorder: A meta-analysis of resting-state functional connectivity. JAMA Psychiatry. 2015;72(6):603–611. doi:10.1001/jamapsychiatry.2015.0071

13. Lynch CJ, Elbau IG, Ng T, et al. Frontostriatal Salience Network Expansion in Individuals in Depression. Vol 633. Springer US; 2024. doi:10.1038/s41586-024-07805-2

14. Nordanskog P, Dahlstrand U, Larsson MR, Larsson EM, Knutsson L, Johanson A. Increase in hippocampal volume after electroconvulsive therapy in patients with depression: a volumetric magnetic resonance imaging study. J ECT. 2010;26(1):62–67. doi:10.1097/YCT.0b013e3181a95da8

15. Takamiya A, Chung JK, Liang KC, Graff-Guerrero A, Mimura M, Kishimoto T. Effect of electroconvulsive therapy on hippocampal and amygdala volumes: Systematic review and meta-analysis. Br J Psychiatry. 2018;212(1):19–26. doi:10.1192/bjp.2017.11

16. Takamiya A, Plitman E, Chung JK, et al. Acute and long-term effects of electroconvulsive therapy on human dentate gyrus. Neuropsychopharmacology. 2019;44(10):1805–1811. doi:10.1038/s41386-019-0312-0

17. Oltedal L, Narr KL, Abbott C, et al. Volume of the Human Hippocampus and Clinical Response Following Electroconvulsive Therapy. Biol Psychiatry. 2018;84(8):574–581. doi:10.1016/j.biopsych.2018.05.017

18. Ousdal OT, Argyelan M, Narr KL, et al. Brain Changes Induced by Electroconvulsive Therapy Are Broadly Distributed. Biol Psychiatry. 2020;87(5):451–461. doi:10.1016/j.biopsych.2019.07.010

19. Argyelan M, Oltedal L, Deng Z De, et al. Electric field causes volumetric changes in the human brain. Elife. 2019;8:1–20. doi:10.7554/eLife.49115

20. Deng Z De, Argyelan M, Miller J, et al. Electroconvulsive therapy, electric field, neuroplasticity, and clinical outcomes. Mol Psychiatry. 2022;27(3):1676–1682. doi:10.1038/s41380-021-01380-y

21. Takamiya A, Bouckaert F, Laroy M, et al. Biophysical mechanisms of electroconvulsive therapy-induced volume expansion in the medial temporal lobe: A longitudinal in vivo human imaging study. Brain Stimul. 2021;14(4):1038–1047. doi:10.1016/j.brs.2021.06.011

22. Madsen TM, Treschow A, Bengzon J, Bolwig TG, Lindvall O, Tingström A. Increased neurogenesis in a model of electroconvulsive therapy. Biol Psychiatry. 2000;47(12):1043–1049. doi:10.1016/S0006-3223(00)00228-6

23. Perera TD, Coplan JD, Lisanby SH, et al. Antidepressant-induced neurogenesis in the hippocampus of adult nonhuman primates. J Neurosci. 2007;27(18):4894–4901. doi:10.1523/JNEUROSCI.0237-07.2007

24. Abe Y, Yokoyama K, Kato T, Yagishita S, Tanaka KF, Takamiya A. Neurogenesis-independent mechanisms of MRI-detectable hippocampal volume increase following electroconvulsive stimulation. Neuropsychopharmacology. 2024;49(8):1236–1245. doi:10.1038/s41386-023-01791-1

25. Andrade C, Bolwig TG. Electroconvulsive therapy, hypertensive surge, blood-brain barrier breach, and amnesia: Exploring the evidence for a connection. J ECT. 2014;30(2):160–164. doi:10.1097/YCT.0000000000000133

26. Diehl DJ, Keshavan MS, Kanal E, Nebes RD, Nichols TE, Gillen JS. Post-ECT increases in MRI regional T2 relaxation times and their relationship to cognitive side effects: A pilot study. Psychiatry Res. 1994;54(2):177–184. doi:10.1016/0165-1781(94)90005-1

27. Kunigiri G, Jayakumar PN, Janakiramaiah N, Gangadhar BN. MRI T(2) relaxometry of brain regions and cognitive dysfunction following electroconvulsive therapy. Indian J Psychiatry. 2007;49(3):195–199.

28. White NS, Leergaard TB, D’Arceuil H, Bjaalie JG, Dale AM. Probing tissue microstructure with restriction spectrum imaging: Histological and theoretical validation. Hum Brain Mapp. 2013;34(2):327–346. doi:10.1002/hbm.21454

29. Loi RQ, Leyden KM, Balachandra A, et al. Restriction spectrum imaging reveals decreased neurite density in patients with temporal lobe epilepsy. Epilepsia. 2016;57(11):1897–1906. doi:10.1111/epi.13570

30. Oltedal L, Kessler U, Ersland L, et al. Effects of ECT in treatment of depression: Study protocol for a prospective neuroradiological study of acute and longitudinal effects on brain structure and function. BMC Psychiatry. 2015;15(1). doi:10.1186/s12888-015-0477-y

31. Madhyastha T, Mérillat S, Hirsiger S, et al. Longitudinal reliability of tract-based spatial statistics in diffusion tensor imaging. Hum Brain Mapp. 2014;35(9):4544–4555. doi:10.1002/hbm.22493

32. Engvig A, Fjell AM, Westlye LT, et al. Memory training impacts short-term changes in aging white matter: A Longitudinal Diffusion Tensor Imaging Study. Hum Brain Mapp. 2012;33(10):2390–2406. doi:10.1002/hbm.21370

33. Schwarz CG, Reid RI, Gunter JL, et al. Improved DTI registration allows voxel-based analysis that outperforms Tract-Based Spatial Statistics. Neuroimage. 2014;94:65–78. doi:10.1016/j.neuroimage.2014.03.026

34. Tustison NJ, Avants BB, Cook PA, et al. Logical circularity in voxel-based analysis: Normalization strategy may induce statistical bias. Hum Brain Mapp. 2014;35(3):745–759. doi:10.1002/hbm.22211

35. Reas ET, Hagler DJ, White NS, et al. Sensitivity of restriction spectrum imaging to memory and neuropathology in Alzheimer’s disease. Alzheimer’s Res Ther. 2017;9(1):1–12. doi:10.1186/s13195-017-0281-7

36. Smith SM, Nichols TE. Threshold-free cluster enhancement: Addressing problems of smoothing, threshold dependence and localisation in cluster inference. Neuroimage. 2009;44(1):83–98. doi:10.1016/j.neuroimage.2008.03.061

37. Fan CC, Loughnan R, Makowski C, et al. Multivariate genome-wide association study on tissue-sensitive diffusion metrics highlights pathways that shape the human brain. Nat Commun. 2022;13(1). doi:10.1038/s41467-022-30110-3

38. Elman JA, Puckett OK, Hagler DJ, et al. Associations between MRI-assessed locus coeruleus integrity and cortical gray matter microstructure. Cereb Cortex. 2022;32(19):4191–4203. doi:10.1093/cercor/bhab475

39. Martin DM, Gálvez V, Loo CK. Predicting retrograde autobiographical memory changes following electroconvulsive therapy: Relationships between individual, treatment, and early clinical factors. Int J Neuropsychopharmacol. 2015;18(12):1–8. doi:10.1093/ijnp/pyv067

40. Ashburner J, C H, R F, I J, C P, Friston KJ. Identifying global anatomical differences: deformation-based morphometry. Hum Brain Mapp. 1998;6:348–57.

41. Nuninga JO, Mandl RCW, Froeling M, et al. Vasogenic edema versus neuroplasticity as neural correlates of hippocampal volume increase following electroconvulsive therapy. Brain Stimul. 2020;13(4):1080–1086. doi:10.1016/j.brs.2020.04.017

42. Dwork AJ, Arango V, Underwood M, et al. Absence of Histological Lesions in Primate Models of ECT and Magnetic Seizure Therapy. Am J Psychiatry. 2004;161:576–578.

43. Sigström R, Göteson A, Joas E, et al. Blood biomarkers of neuronal injury and astrocytic reactivity in electroconvulsive therapy. Mol Psychiatry. 2024;(March):1–9. doi:10.1038/s41380-024-02774-4

44. Cudna A, Bronisz E, Jopowicz A, Kurkowska-Jastrzębska I. Changes in serum blood-brain barrier markers after bilateral tonic-clonic seizures. Seizure. 2023;106(February 2022):129–137. doi:10.1016/j.seizure.2023.02.012

45. Zimmermann R, Schmitt H, Rotter A, Sperling W, Kornhuber J, Lewczuk P. Transient increase of plasma concentrations of amyloid β peptides after electroconvulsive therapy. Brain Stimul. 2012;5(1):25–29. doi:10.1016/j.brs.2011.01.007

46. Rush G, O’Donovan A, Nagle L, et al. Alteration of immune markers in a group of melancholic depressed patients and their response to electroconvulsive therapy. J Affect Disord. 2016;205:60–68. doi:10.1016/j.jad.2016.06.035

47. Lehtimäki K, Keränen T, Huuhka M, et al. Increase in plasma proinflammatory cytokines after electroconvulsive therapy in patients with depressive disorder. J ECT. 2008;24(1):88–91. doi:10.1097/YCT.0b013e3181571abb

48. Montagne A, Barnes SR, Sweeney MD, et al. Blood-Brain barrier breakdown in the aging human hippocampus. Neuron. 2015;85(2):296–302. doi:10.1016/j.neuron.2014.12.032

49. Watts B V., Peltzman T, Shiner B. Mortality after electroconvulsive therapy. Br J Psychiatry. 2021;219(5):588–593. doi:10.1192/bjp.2021.63

50. Rhee TG, Sint K, Olfson M, Gerhard T, H Busch S, Wilkinson ST. Association of ECT With Risks of All-Cause Mortality and Suicide in Older Medicare Patients. Am J Psychiatry. 2021;178(12):1089–1097. doi:10.1176/appi.ajp.2021.21040351

51. Tørring N, Sanghani SN, Petrides G, Kellner CH, Østergaard SD. The mortality rate of electroconvulsive therapy: a systematic review and pooled analysis. Acta Psychiatr Scand. 2017;135(5):388–397. doi:10.1111/acps.12721

52. Osler M, Rozing MP, Christensen GT, Andersen PK, Jørgensen MB. Electroconvulsive therapy and risk of dementia in patients with affective disorders: a cohort study. The Lancet Psychiatry. 2018;5(4):348–356. doi:10.1016/S2215-0366(18)30056-7

53. Hjerrild S, Kahlert J, Buchholtz PE, Rosenberg R, Videbech P. Long-term risk of developing dementia after electroconvulsive therapy for affective disorders. J ECT. 2021;37(4):250–255. doi:10.1097/YCT.0000000000000770

54. Zeiler FA, Matuszczak M, Teitelbaum J, Gillman LM, Kazina CJ. Electroconvulsive therapy for refractory status epilepticus: A systematic review. Seizure. 2016;35:23–32. doi:10.1016/j.seizure.2015.12.015

55. Erchinger VJ, Evjenth Sørhaug OJ, Aukland SM, et al. Effects of Electroconvulsive Therapy on Brain Structure - a Neuroradiological Investigation into White Matter Hyperintensities, Atrophy, and Microbleeds. Biol Psychiatry Cogn Neurosci Neuroimaging 2024 doi: 10.1016/j.bpsc.2024.12.004.

56. Leaver AM, Espinoza R, Pirnia T, Joshi SH, Woods RP, Narr KL. Modulation of Intrinsic Brain Activity by Electroconvulsive Therapy in Major Depression. Biol Psychiatry Cogn Neurosci Neuroimaging. 2016;1(1):77–86. doi:10.1016/j.bpsc.2015.09.001

57. Argyelan M, Lencz T, Kaliora S, et al. Subgenual cingulate cortical activity predicts the efficacy of electroconvulsive therapy. Transl Psychiatry. 2016;6(4):e789–6. doi:10.1038/tp.2016.54

58. Takamiya A, Kishimoto T, Hirano J, et al. Neuronal network mechanisms associated with depressive symptom improvement following electroconvulsive therapy. Psychol Med. 2021;51(16):2856–2863. doi:10.1017/S0033291720001518

59. Mattson MP and Leak RK. The hormesis principle of neuroplasticity and neuroprotection. Cell Metab 2024; 36(2): 315–337.

