## SupplementaryMaterial for "Seizure-induced Transient Disruptive Changes in Brain Microstructure"

### Supplementary Material

Supplementary Methods

Supplementary Figure 1. Results of the whole-brain voxel-wise analysis of mean diffusivity.

Supplementary Figure 2. Effect sizes of the impact of a single seizure

Supplementary Figure 3. Results of the voxel-wise analysis restricted to the white matter.

Supplementary Figure 4. Results of the voxel-wise analysis restricted to the gray matter.

Supplementary Figure 5. Correlations between post-ictal reorientation time and changes in RSI-metrics.

Supplementary Figure 6. Short- and long-term changes in RSI-metrics.

Supplementary Figure 7. Ventricular volumetric change following a single seizure.

Supplementary References

**Supplementary Methods**

*ECT*

ECT was administered three times a week with a Thymatron Sytem IV (Somatics LLC, Venice, FL, USA). All participants received brief pulse (0.5 ms) right unilateral (RUL) ECT. The initial stimulus charge was determined by age (i.e., [Individual's age in years] % Energy dial of the Thymatron device)[1]. Thiopental or propofol and succinylcholine were used for anesthesia and muscle relaxant, respectively. Duration of electroencephalographic (EEG) seizure was measured by two-channel EEG (Fp1-M1, Fp2-M2, international 10-20 system).

*Image acquisition and processing*

High-resolution 3D T1-weighted images (TE/TR = 2.9/6.7 ms; inversion time = 600 ms; flip angle = 8 degree; FOV = 256 mm; voxel size = 1.0 × 1.0 × 1.0 $mm^3$) and multi-shell diffusion MRI with a single-shot pulsed-field gradient spin-echo EPI sequence, an optimized sequence for RSI (TE/TR = 85/7000 ms; FOV = 240 mm; matrix = 96 × 96 × 55 with 4 b-values [b=0, 500, 1000, and 4000 s/$mm^2$ and 6, 6, and 15 unique directions for the nonzero b-values, respectively] ), were acquired on a 3T Discovery MR750 system with a 32-channel head coil (GE HealthCare, Waukesha, WI).

T1-weighted structural images were processed using the longitudinal pipeline of the Computational Anatomy Toolbox (CAT12; version 12.9). CAT12 was used 1) to explore ventricular/CSF changes following one session of ECT, 2) creating brain masks in whole-brain analyses, and 3) to estimate precise registration parameters in diffusion-weighted imaging analysis. Ventricular volumes were calculated based on the Hammers Atlas implemented in CAT12. This atlas was selected because the temporal horn of the lateral ventricles and lateral ventricles excluding temporal horn are labelled separately. Right and left volumes of the lateral ventricles were summed. Those ventricular volumes were normalized by regressing out the total intracranial volume (TIV), which was calculated using CAT12. Second, to create brain masks in MNI152NLin2009cAsym space, the T1 template implemented in CAT12 was segmented into gray matter (GM), white matter (WM), and cerebrospinal fluid (CSF). The segmented tissue maps were thresholded at 0.5 and binarized. For brain tissue mask, GM and WM masks were combined, followed by subtraction of CSF mask to exclude potential overlapped regions. Additionally, brain masks including one tissue type (i.e., GM, WM, or CSF) were also created by subtracting the other two tissue types. Because the cerebellum and lower brain stem were not completely covered in all scans, these regions were excluded from the analyses. Third, for better registration in DWI analysis, intermediate 3D T1files using the CAT12 longitudinal pipeline were extracted. In brief, the CAT12 pipeline first realigns the images from two time points using inverse-consistent rigid-body registrations with intra-subject bias-field corrections, and it creates subject-specific average images between the time points (https://neuro-jena.github.io/cat12-help/#long_opt). The realigned images and average images were utilized for the registration steps in the following diffusion-weighted imaging analysis.

Diffusion-weighted images were first visually checked for quality and artifacts. Images were pre-processed using an in-house pipeline, including correction for B0 distortion, motion and eddy current distortions, gradient nonlinearity distortions, and head motion. Diffusion-weighted images were applied a one voxel median filter to improve registration in longitudinal analysis pipeline[2]. B0 images were registered to skull-stripped T1-weighted images using Advanced Normalization Tools (ANTs) (https://github.com/ANTsX/ANTs), and then the resultant transformation matrices were applied to pre-processed diffusion-weighted images. Visual inspection of the pre-processed images was conducted and images from one participant were excluded because of motion artifact.

To optimally align diffusion-weighted images within each individual subject between two time points, study-specific T1-weighted templates in each group were created using the subject-specific average images with antsMultivariateTemplateConstruction2.sh, and then all images were warped to the study-specific template space[2–5]. Finally, all images were warped to the common standard MNI152NLin2009cAsym space and were smoothed with a 2-mm full-width at half-maximum Gaussian kernel. In all the registration steps, transformation matrices calculated using T1-weighted images with antsRegistrationSyN.sh[4,5]were applied to corresponding diffusion-weighted images using antsApplyTransforms.

RSI model was fit using an in-house Matlab (the MathWorks, Natick, MA)-based script developed by the UCSD Multimodal Imaging Laboratory, which has been applied to other datasets by other groups[6,7]. Theoretical background and technical details are presented elsewhere[8]. Measures derived from the RSI model fit include restricted, hindered, and isotropic diffusion. These RSI maps were normalized by taking the norm (square root of the sum of squares) of each fractional map and dividing by the norm of all fractional maps, making them unitless with range between 0 to 1. As an additional analysis, a diffusion tensor model was also fit to the data with b=1000 using FSL to generate mean diffusivity maps (Supplementary Figure 1).

To investigate the short- (<2 weeks) and long-term (6 months) trajectories of the identified RSI changes in the main analysis, we analyzed follow-up data collected at TP3 (timepoint 3: 7–14 days post-ECT; n=23) and TP4 (timepoint 4: 6 months post-ECT; n=17). Because the primary aim of this study was to investigate the effects of a single seizure on the human brain, this follow-up analysis was limited to the RSI changes identified in the whole-brain analysis between TP1 and TP2 (Figure 1) to determine whether these changes were transient. Due to missing data at TP3 and TP4, a linear mixed effects model was employed, with age and sex included as covariates. Although our primary objective of this analysis was to assess whether the observed changes at TP2 returned to baseline levels, we adhered to the standard reporting approach for linear mixed-effects models (i.e., the main effect of time was reported first, followed by pairwise comparisons). Raw p-values are reported, while results with Bonferroni correction for multiple comparisons are presented in the Supplementary Figure 6.

**Supplementary Figure 1. Results of the whole-brain voxel-wise analysis of mean diffusivity.** (**A**) A single seizure increased mean diffusivity across widely distributed brain regions bilaterally. Red indicates regions with increased mean diffusivity, while blue represents regions with decreased mean diffusivity following a single seizure. Only brain regions with increased mean diffusivity were identified at the statistical threshold of family-wise error (FWE)-corrected p <0.05, determined using threshold-free cluster enhancement (TFCE). (**B**) Extracted values of mean diffusivity in the regions identified in the ECT group showed no significant changes in the healthy control group (t=1.5, p=0.16) or the electrical cardioversion group (t=1.8, p=0.09). These boxplots are presented for illustrative purposes. The unit for mean diffusivity is $mm^2$/s.

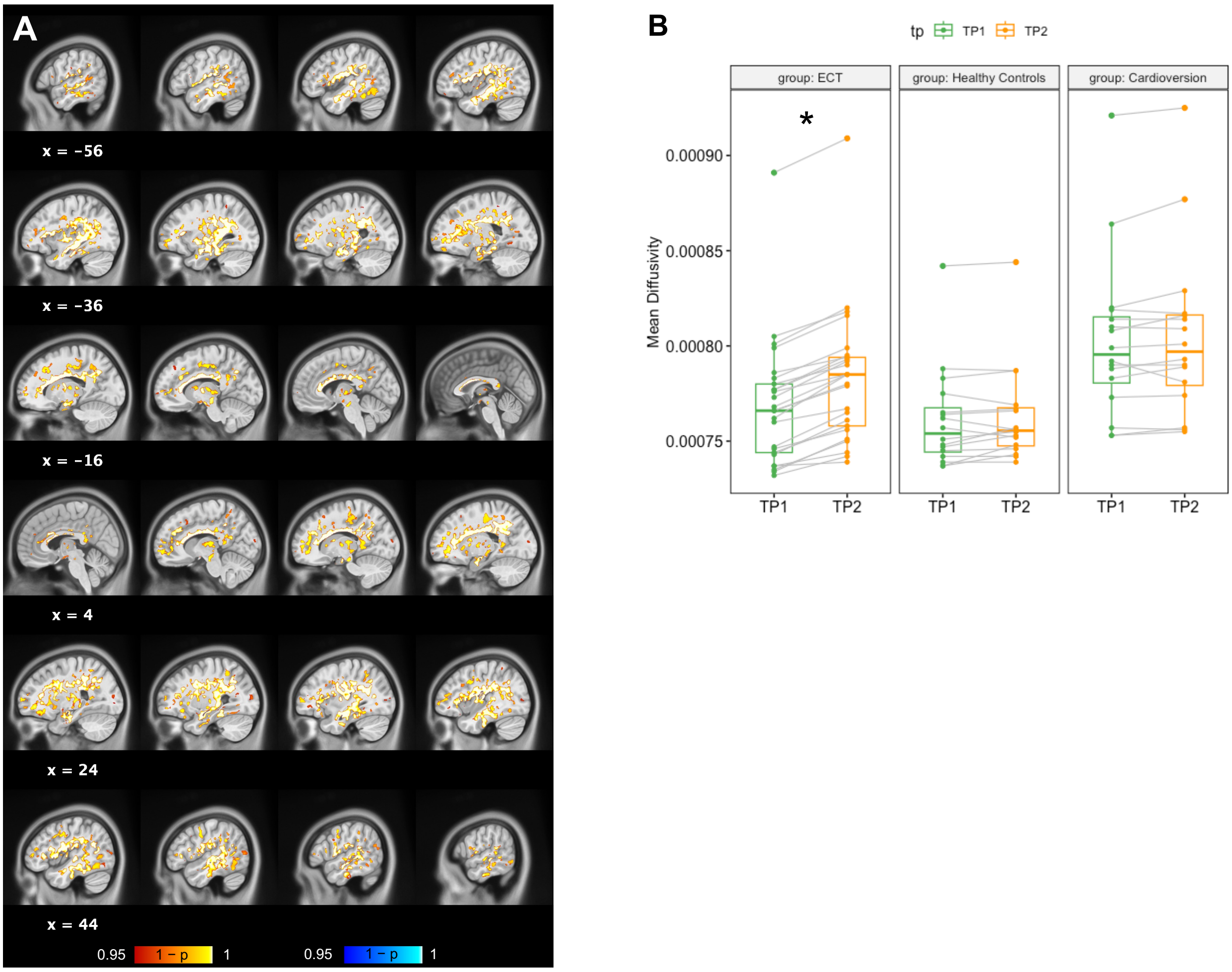

**Supplementary Figure 2. Effect sizes of the impact of a single seizure on free water (A), restricted diffusion (B), hindered diffusion (C), and mean diffusivity (D).** Cohen's d was calculated using t-values, and effect sizes of d >0.3 are displayed in the figures. Red indicated increases, while blue indicates decreases in each metric following a single seizure. Although no significant brain regions for hindered diffusion were identified in the statistical analysis reported in main text, the effect size map reveals a regional pattern of increases and decreases (**C**).

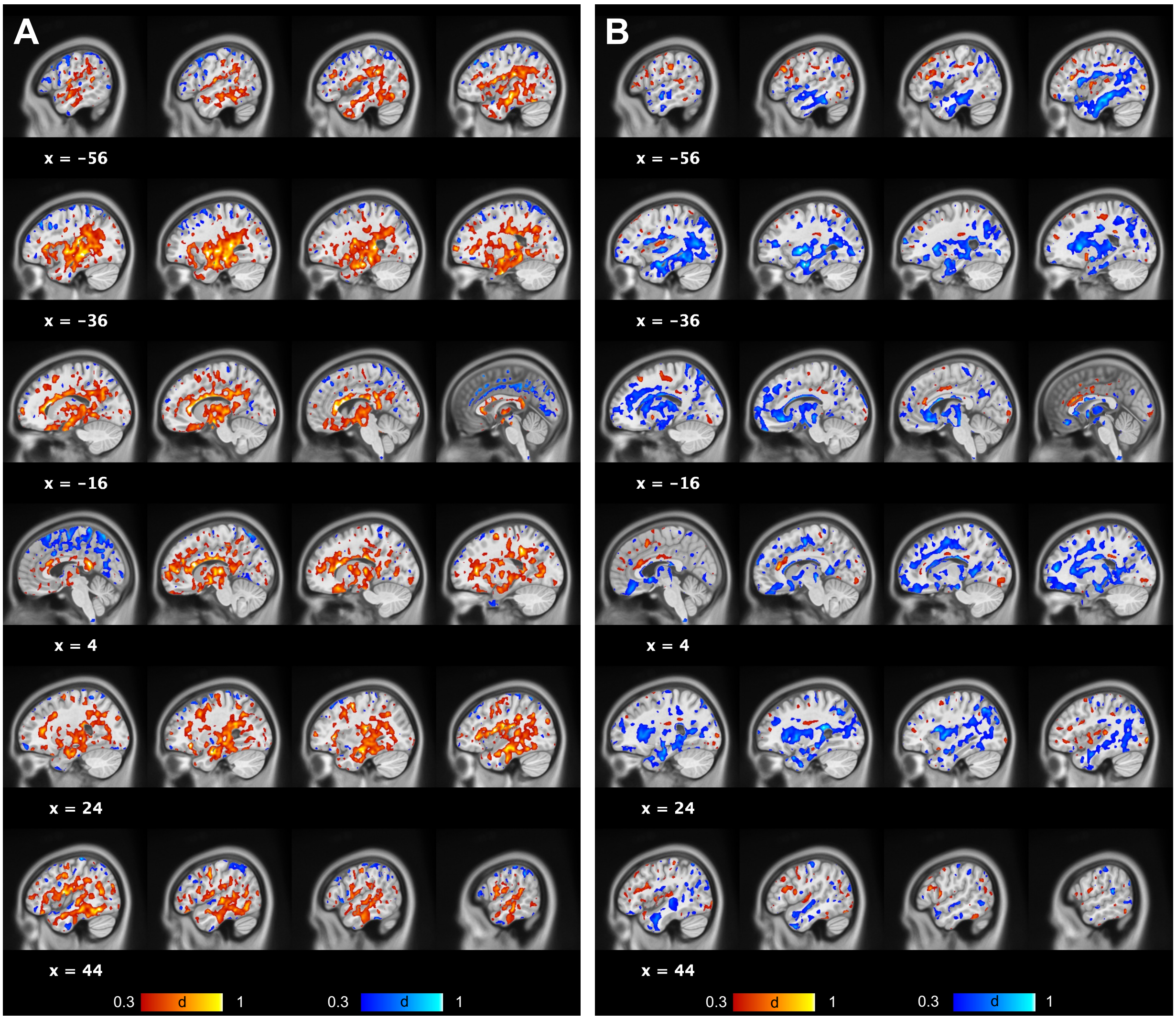

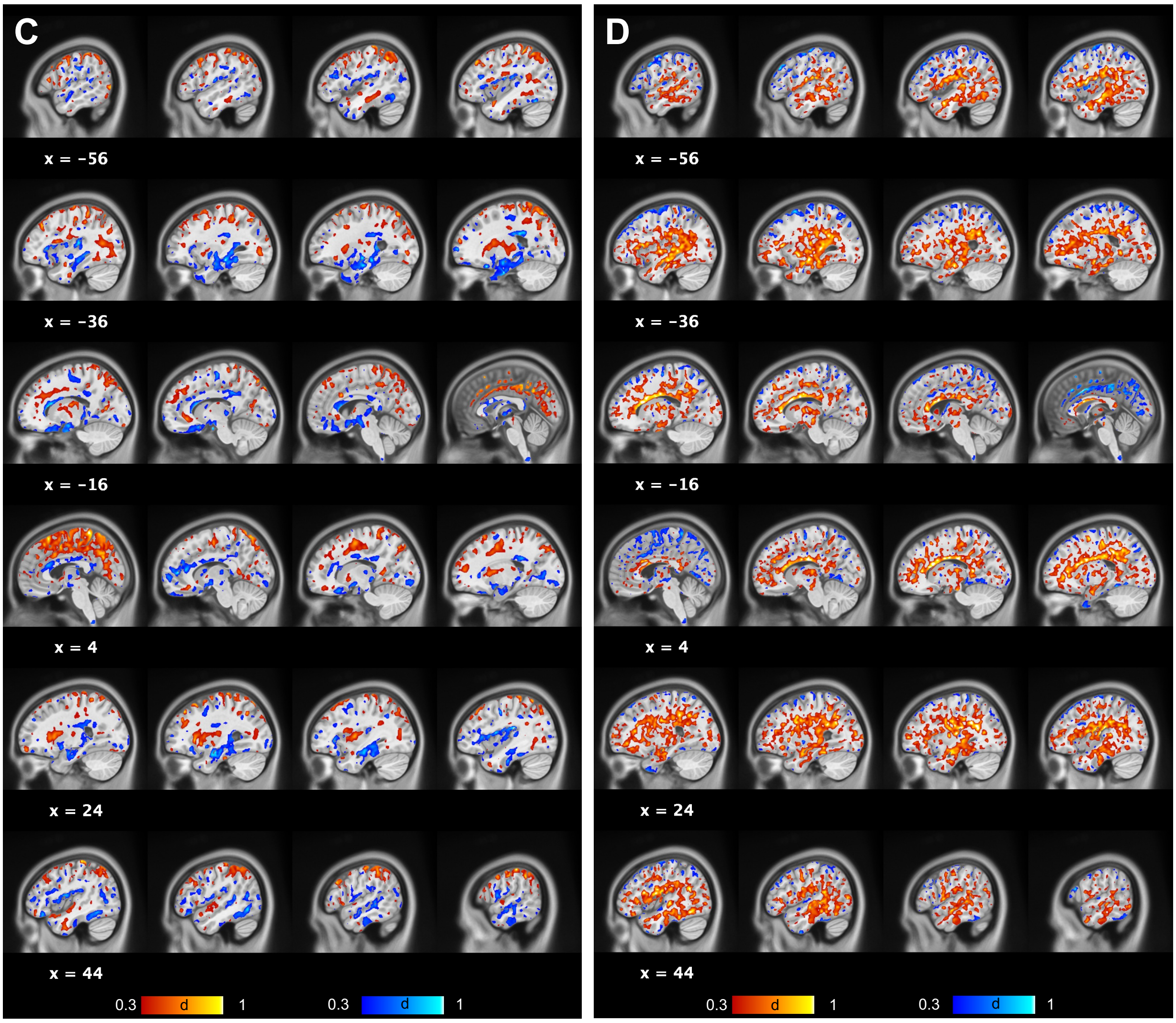
C
x = -56
x = -36
x = -16
x = 4
x = 24
x = 44
0.3 d 1
0.3 d 1
D
x = -56
x = -36
x = -16
x = 4
x = 24
x = 44
0.3 d 1
0.3 d 1

**Supplementary Figure 3. Results of the voxel-wise analysis restricted to the white matter.** (**A**) Increased free water was observed in the periventricular white matter, including the corpus callosum. (**B**) Decreased restricted diffusion was primarily identified in the corpus callosum and internal/external capsule. (**C**) Increased mean diffusivity was identified across widely distributed brain regions bilaterally, extending beyond the periventricular regions. Red indicates increases, while blue indicates decreases following a single seizure. The statistical threshold was set at FWE-corrected p <0.05, determined using TFCE.

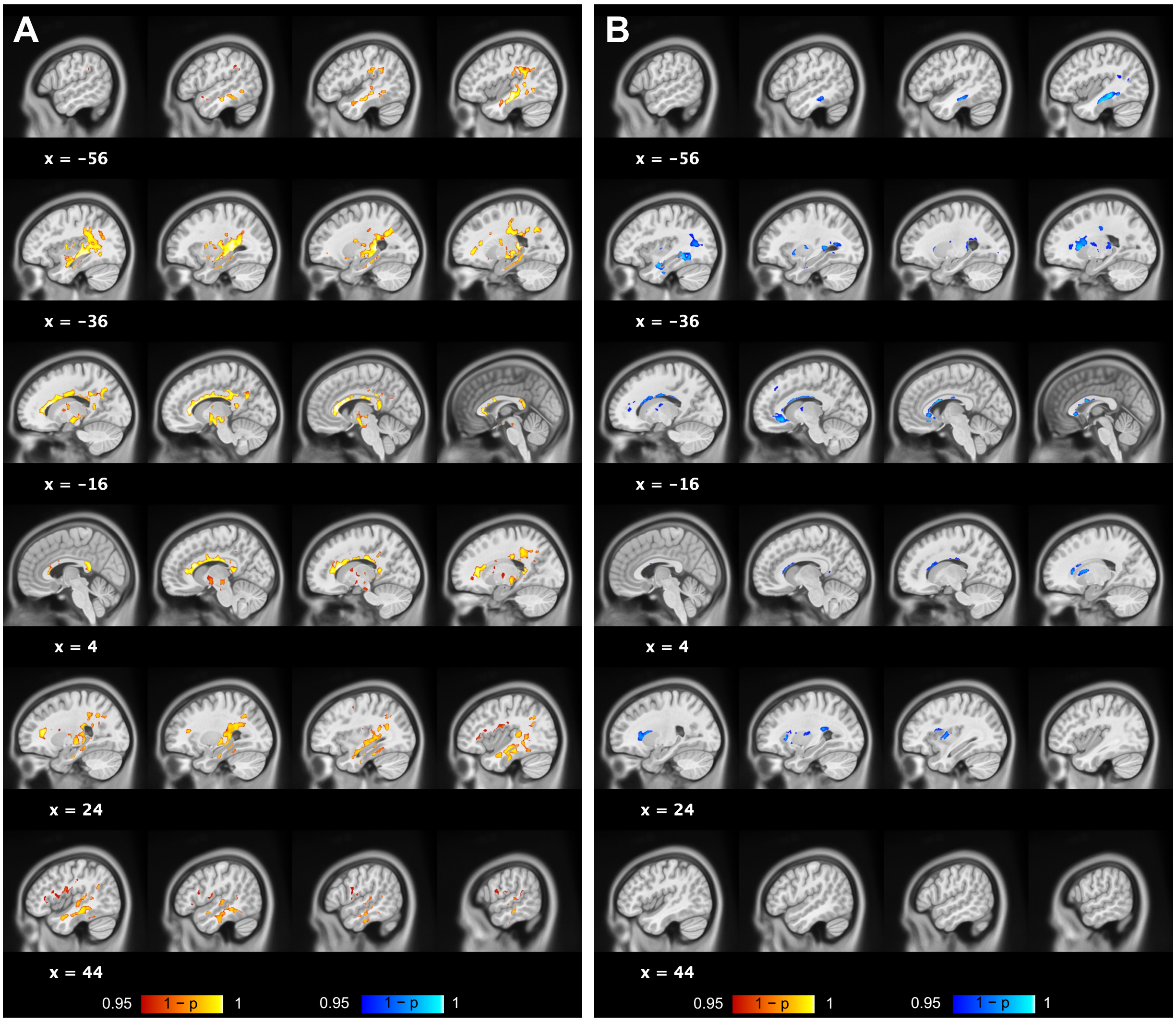

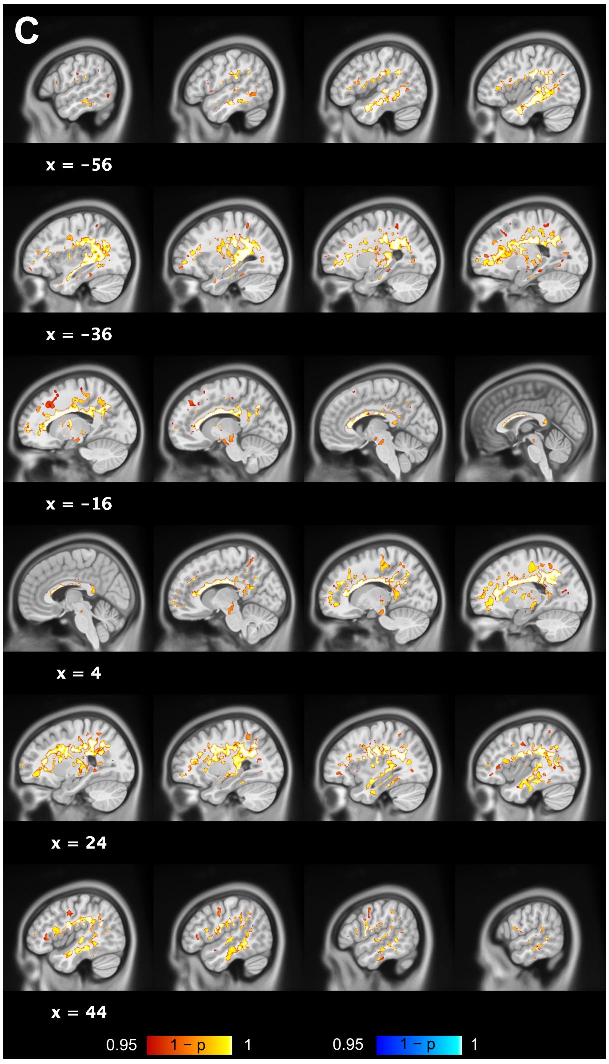
C
x = −56
x = −36
x = −16
x = 4
x = 24
x = 44
0.95
1 − p
1
0.95
1 − p
1

**Supplementary Figure 4. Results of the voxel-wise analysis restricted to the gray matter.** (**A**) Increased free water was observed in the hippocampus, thalamus, putamen, claustrum, insula, and orbitofrontal cortex. (**B**) Decreased restricted diffusion was identified in the hippocampus, putamen, and claustrum, predominantly in the left hemisphere. (**C**) The gray matter analysis revealed increased hindered diffusion in regions adjacent to the longitudinal fissure and decreased hindered diffusion in the hippocampus. (**D**) Increased mean diffusivity was identified in the hippocampus and insula. Red indicates increase, while blue indicates decreases following a single seizure. The statistical threshold was set at FWE-corrected p <0.05, determined using TFCE.

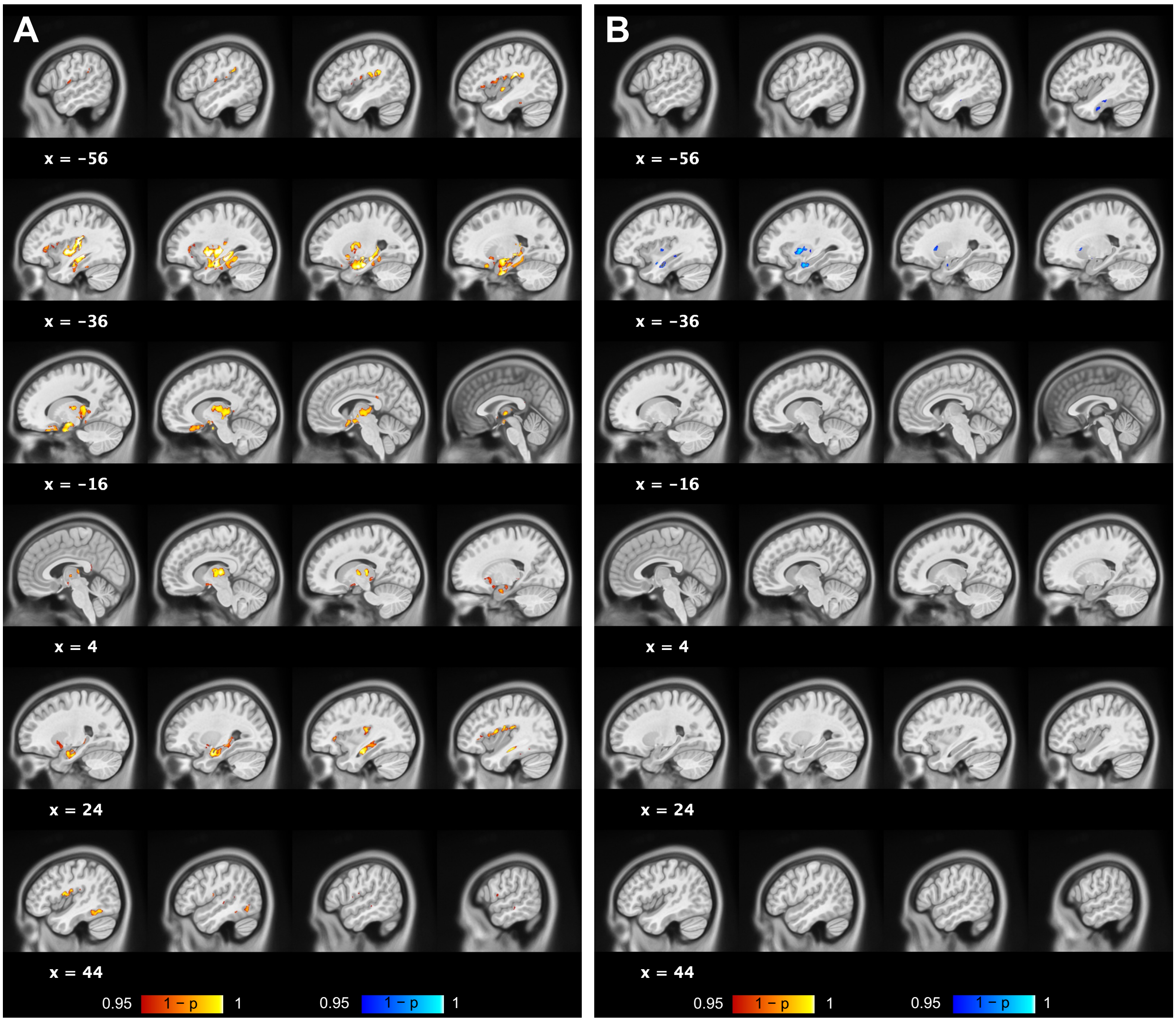

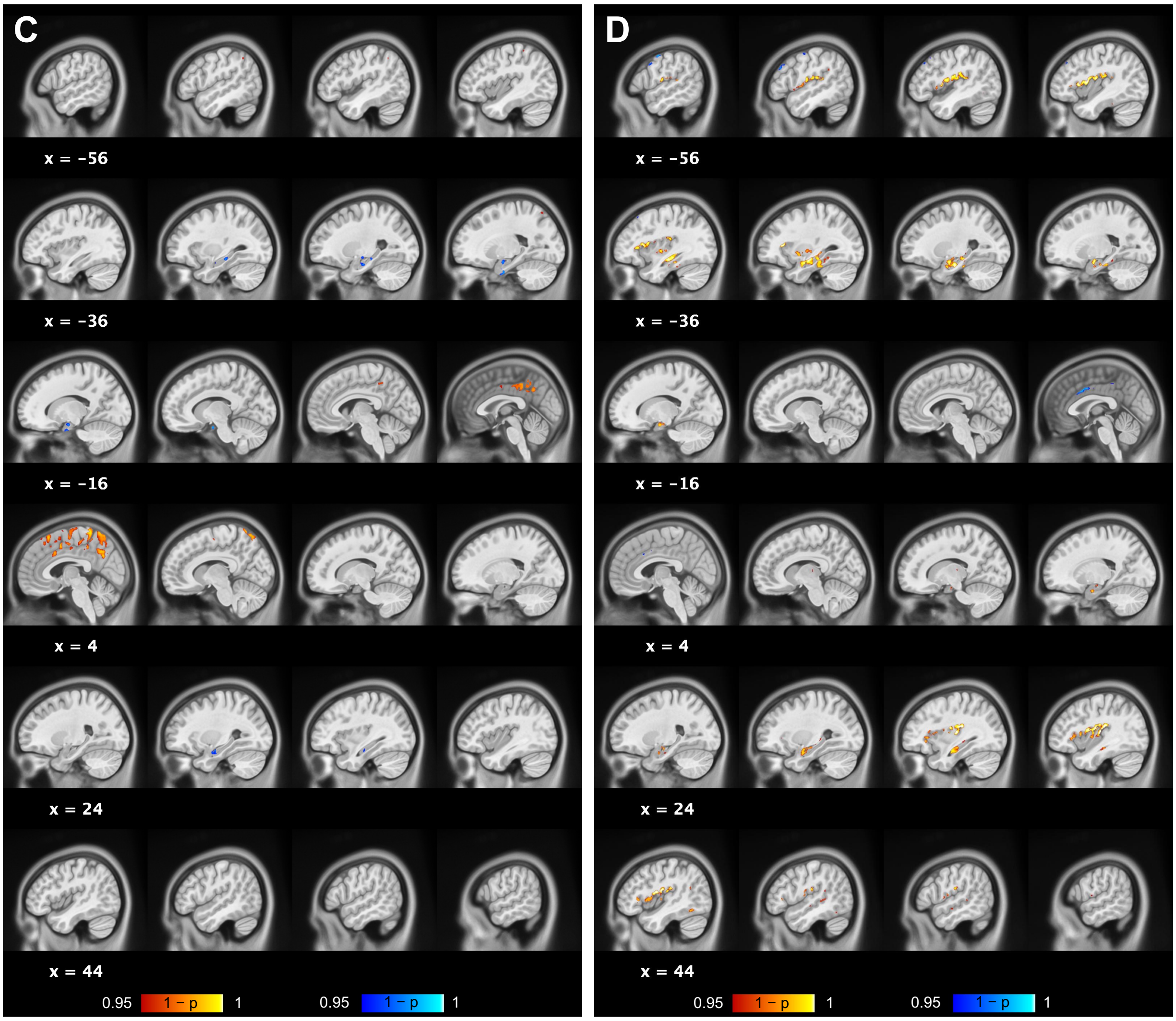
C
x = −56
x = −36
x = −16
x = 4
x = 24
x = 44
0.95
1 − p
1
0.95
1 − p
1
D
x = −56
x = −36
x = −16
x = 4
x = 24
x = 44
0.95
1 − p
1
0.95
1 − p
1

**Supplementary Figure 5. Correlations between post-ictal reorientation time and changes in RSI-metrics.** Post-ictal reorientation time was not associated with changes in free water (**A**: r=0.11, df=23, p=0.92) or restricted diffusion (**B**: r=0.10, df=23, p=0.64).

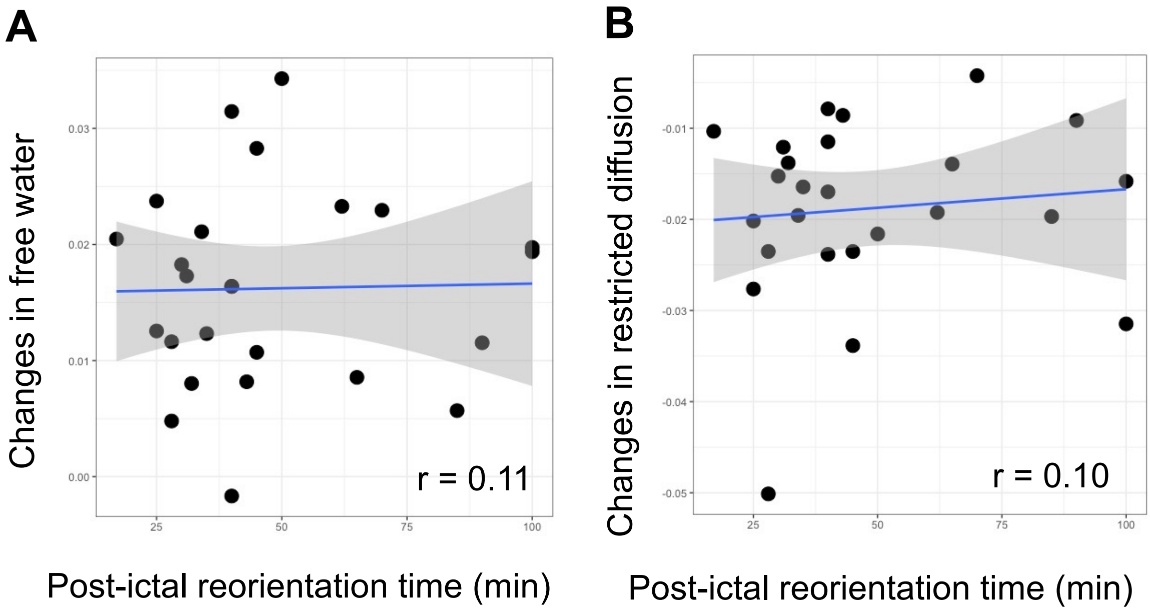

**Supplementary Figure 6. Short- and long-term changes in RSI-metrics.** Linear mixed-effects models showed a significant effect of time on free water (**A**: $F_{3,62.5} = 12.9$, p<0.001) and restricted diffusion (**B**: $F_{3,62.2} = 21.8$, p<0.001). (**A**) Post-hoc pairwise comparisons revealed significant differences in free water between TP2 and TP3 (t=–4.8, p<0.001), while there was no significant difference between TP1 and TP3 (t=0.69, p=0.50), suggesting that the increases in free water following a single seizure returned to baseline levels by TP3. (**B**) Post-hoc pairwise comparisons revealed significant differences in restricted diffusion between TP2 and TP4 (t=5.5, p<0.001), and TP1 and TP3 (t=–4.7, p<0.001), while there was no difference between TP1 and TP4 (t=–0.90, p=0.37). These results suggest that the decrease in restricted diffusion returned to baseline levels by TP4 but not by TP3. In summary, the observed changes following a single seizure (i.e., increased free water and decreased restricted diffusion at TP2) were transient and reversible. Double asterisks (**) indicate statistical significance at p <0.05 with Bonferroni correction for multiple comparisons.

TP1: 2 hours before the first ECT session (i.e., around 8 am); TP2: 2 hours after the first ECT session (i.e., around 12 pm); TP3: 7–14 days after the last ECT session; TP4: 6 months after the last ECT session

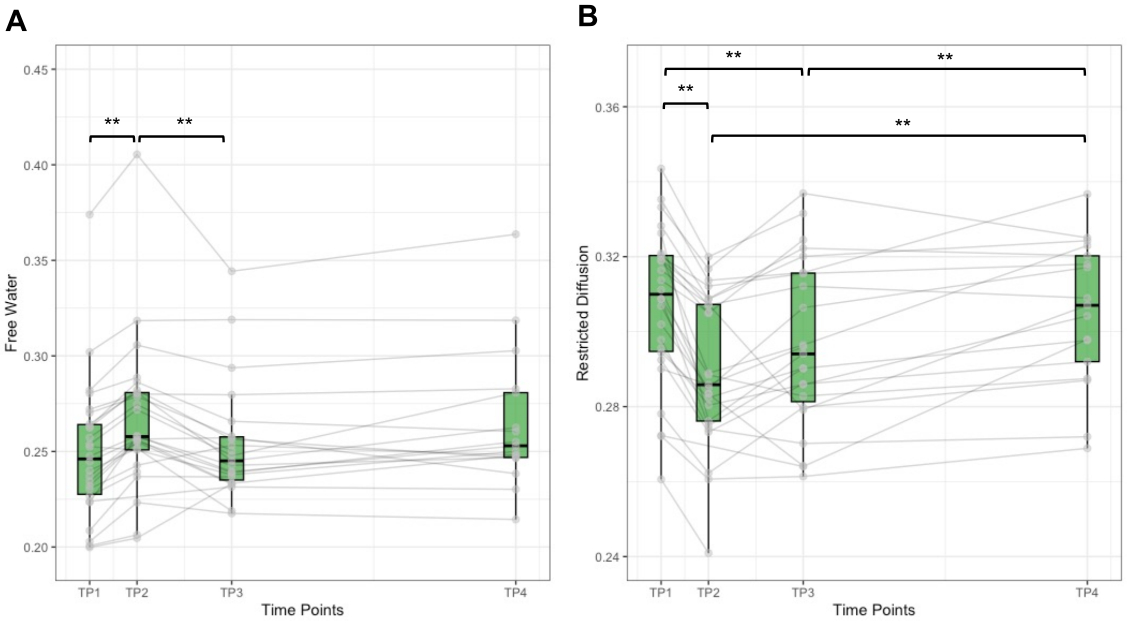

**Supplementary Figure 7. Ventricular volume change following a single seizure.** (**A**) Deformation-based morphometry revealed significant volume expansion in the CSF regions following a single seizure. The statistical threshold was set at FWE-corrected p <0.05, determined using TFCE. Red indicates volume expansion. Only brain regions showing volume expansion were identified. (**B**) Linear mixed effect models, including age and sex as covariates, revealed a significant group by time interaction for volumes of the temporal horn of the lateral ventricle defined by the Hammers Atlas implemented in the CAT12 ($F_{2,54}$=5.75, p=0.005). Post-hoc within-subject comparisons found that only the ECT group showed significant increase in the volumes of the temporal horn (t=4.70, p<0.001), while healthy control (HC) (t=0.21, p=0.84) or electrical cardioversion (ECV) group (t=0.07, p=0.94) did not show significant changes. The unit for volumes is $\times 10^3$ $mm^2$. Volumes of the lateral ventricles excluding temporal horn did not show group by time interaction ($F_{2,54}$=1.35, p=0.27). However, post-hoc within-subject comparisons of ECT group showed significant increase in the volumes of the lateral ventricles excluding temporal horn (t=2.9, p=0.005), while HC (t=0.96, p=0.34) or ECV group (t=0.36, p=0.72) did not show significant changes. Volumes of the third ventricles did not show a significant group by time interaction ($F_{2,54}$=0.48, p=0.62), with post-hoc comparisons showing no significant changes in volumes of the third ventricles (ECT: t=0.97, p=0.34; HC: t=0.54, p=0.59; ECV: t=–0.45, p=0.65). These results suggest that single ECT session induced volume expansion of the ventricle, especially the temporal horn of the lateral ventricle. *p<0.05/3=0.017

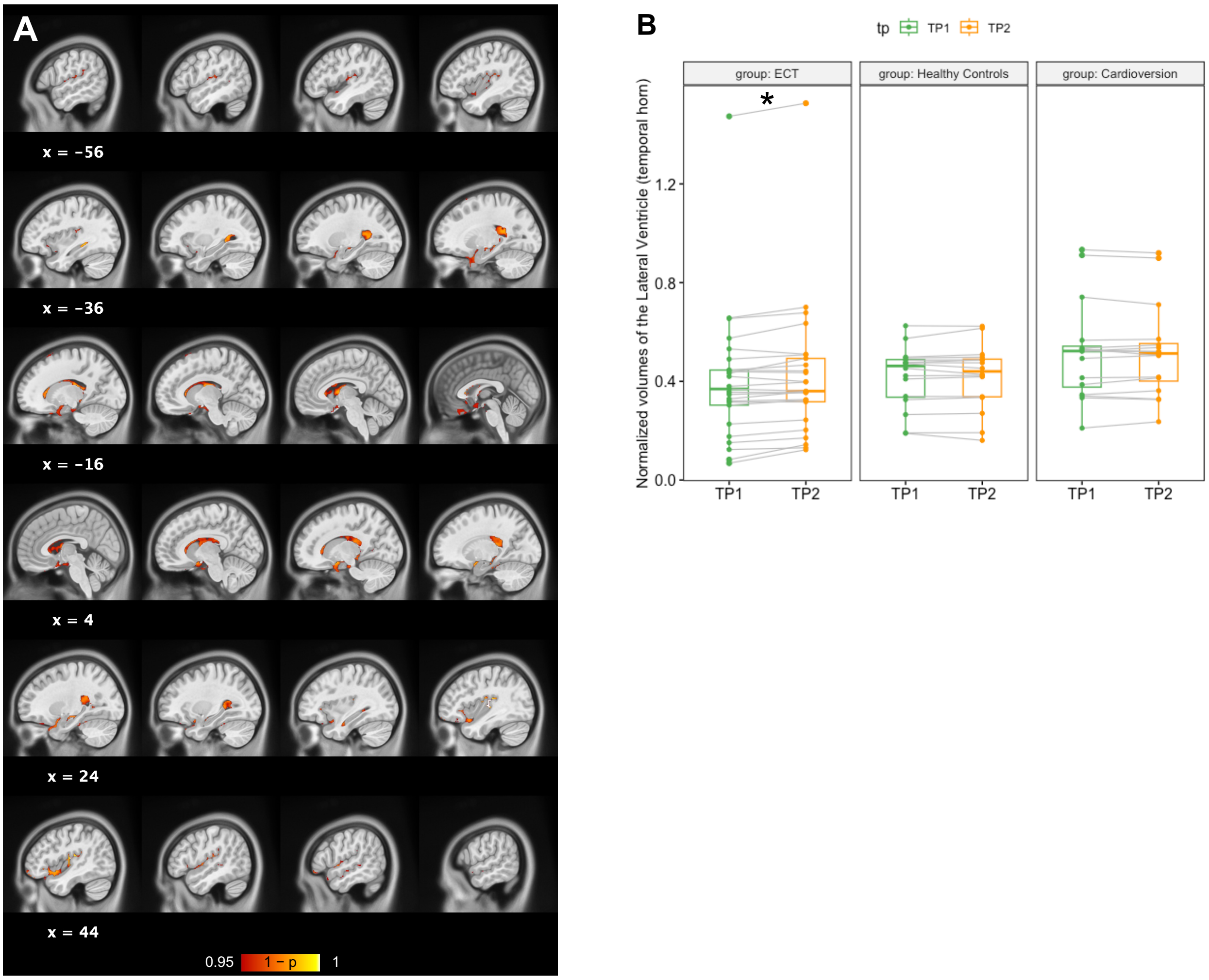

Supplementary References


1. Oltedal L, Kessler U, Ersland L, et al. Effects of ECT in treatment of depression: Study protocol for a prospective neuroradiological study of acute and longitudinal effects on brain structure and function. *BMC Psychiatry*. 2015;15(1). doi:10.1186/s12888-015-0477-y
2. Madhyastha T, Mérillat S, Hirsiger S, et al. Longitudinal reliability of tract-based spatial statistics in diffusion tensor imaging. *Hum Brain Mapp*. 2014;35(9):4544-4555. doi:10.1002/hbm.22493
3. Engvig A, Fjell AM, Westlye LT, et al. Memory training impacts short-term changes in aging white matter: A Longitudinal Diffusion Tensor Imaging Study. *Hum Brain Mapp*. 2012;33(10):2390-2406. doi:10.1002/hbm.21370
4. Schwarz CG, Reid RI, Gunter JL, et al. Improved DTI registration allows voxel-based analysis that outperforms Tract-Based Spatial Statistics. *Neuroimage*. 2014;94:65-78. doi:10.1016/j.neuroimage.2014.03.026
5. Tustison NJ, Avants BB, Cook PA, et al. Logical circularity in voxel-based analysis: Normalization strategy may induce statistical bias. *Hum Brain Mapp*. 2014;35(3):745-759. doi:10.1002/hbm.22211
6. Reas ET, Hagler DJ, White NS, et al. Sensitivity of restriction spectrum imaging to memory and neuropathology in Alzheimer's disease. *Alzheimer's Res Ther*. 2017;9(1):1-12. doi:10.1186/s13195-017-0281-7
7. Loi RQ, Leyden KM, Balachandra A, et al. Restriction spectrum imaging reveals decreased neurite density in patients with temporal lobe epilepsy. *Epilepsia*. 2016;57(11):1897-1906. doi:10.1111/epi.13570
8. White NS, Leergaard TB, D'Arceuil H, Bjaalie JG, Dale AM. Probing tissue microstructure with restriction spectrum imaging: Histological and theoretical validation. *Hum Brain Mapp*. 2013;34(2):327-346. doi:10.1002/hbm.21454
